# Elevated arginine levels in liver tumors promote metabolic reprogramming and tumor growth

**DOI:** 10.1101/2022.04.26.489545

**Authors:** Dirk Mossmann, Sujin Park, Brendan Ryback, Diana Weißenberger, Marco Colombi, Sravanth K. Hindupur, Eva Dazert, Mairene Coto-Llerena, Ercan Caner, Veronica J. Cenzano, Salvatore Piscuoglio, Fatima Bosch, Luigi M. Terracciano, Uwe Sauer, Michael N. Hall

## Abstract

Arginine auxotropy, due to reduced expression of urea cycle genes, is common in cancer. However, little is known about the levels of arginine in these cancers. Here, we report that arginine levels are elevated in hepatocellular carcinoma (HCC) despite reduced expression of urea cycle enzymes. Liver tumors accumulate high levels specifically of arginine via increased uptake and, more importantly, via suppression of arginine-to-polyamine conversion due to reduced arginase 1 (ARG1) and agmatinase (AGMAT) expression. Furthermore, the high levels of arginine are required for tumor growth. Mechanistically, high levels of arginine promote tumorigenesis via transcriptional regulation of metabolic genes, including upregulation of asparagine synthetase (ASNS). ASNS-derived asparagine further enhances arginine uptake, creating a positive feedback loop to sustain high arginine levels and oncogenic metabolism. Thus, arginine is a novel second messenger-like molecule that reprograms metabolism to promote tumor growth.

## Results

### Elevated arginine levels are necessary for liver tumorigenesis

To identify metabolic alterations in HCC, we performed untargeted metabolomics on liver tumors isolated from a previously described mTOR-driven HCC mouse model (*1-3*). In this mouse model, constitutively high mTOR signaling, due to liver-specific double-knockout of the tumor suppressors TSC1 and PTEN (hereafter referred to as L-dKO), drives the development of hepatomegaly, hepatosteatosis, and multiple high-grade HCC within 20 weeks of age (*1, 2*). Using untargeted metabolomics, we detected 6734 and 7629 ions in positive and negative mode, respectively. Allowing relaxed annotation (see Materials and Methods), 3467 metabolites could be tentatively identified of which 916 were significantly altered in abundance, in L-dKO tumors compared to control liver tissues (**fig. S1B**). The metabolic profiles of tumors and control liver tissues separated by principal component analysis (PCA) and hierarchical clustering (**Fig. 1A** and **fig. S1A**). Metabolic pathway enrichment analysis (MPWEA) revealed that amino acid metabolic pathways were commonly altered in L-dKO tumors (**Fig. 1B** and **table S1**). To investigate if amino acid levels were altered in tumors, we performed targeted profiling of amino acids. Interestingly, arginine was elevated in L-dKO tumors while all other amino acids were either unchanged or decreased (**Fig. 1C** and **fig. S1C**). This observation was surprising as liver tumors, and other types of cancer, have been shown to be deficient in arginine synthesis due to suppression of the urea cycle (*4-6*). However, the level of arginine in tumors with alterations in urea cycle enzymes is unknown with the exception of renal cell carcinoma (RCC) where suppression of arginine synthesis results in decreased tumoral arginine levels (*7*).

**Figure 1.**
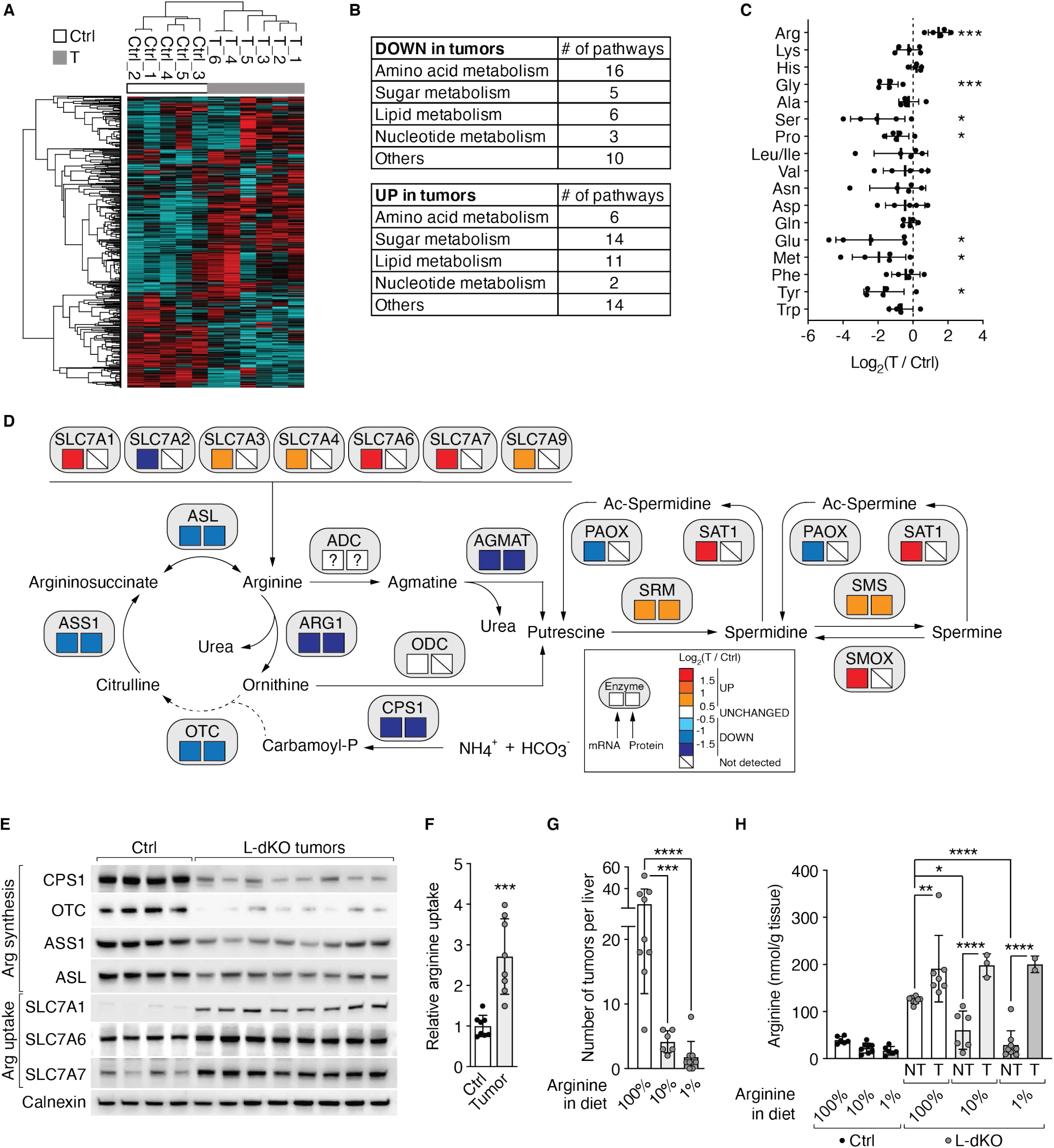
Arginine is elevated in liver tumors and promotes tumor formation. **A**. Hierarchical clustering of significantly deregulated metabolites from control (Ctrl) liver tissues and tumors (T) from liver-specific *Tsc1* and *Pten* double-knockout (hereafter, L-dKO) mice. n=5 (Ctrl) and 6 (L-dKO). **B**. Up-and down-regulated metabolic pathways in L-dKO tumors compared to Ctrl liver tissues after PWEA. **C**. Amino acid profile of L-dKO tumor tissues relative to Ctrl liver tissues (log_2_ ratio). Unpaired t test, *p<0.05, ***p<0.001. n=5 (Ctrl) and 5 (L-dKO). **D**. Schematic representation of arginine and polyamine metabolism. Boxes below enzymes indicate changes in mRNA (left box) and protein (right box) levels in L-dKO tumors compared to Ctrl livers, respectively. Color coding according to level of log_2_-fold change as indicated. “?” indicates unknown identity. n=6 (Ctrl) and 12 (L-dKO). **E**. Immunoblot analyses of arginine synthesizing enzymes (CPS1, OTC, ASS1, ASL) and arginine transporters (SLC7A1, SLC7A6, SLC7A7) in Ctrl liver and L-dKO tumor tissues. Calnexin serves as loading control. n=4 (Ctrl) and 8 (L-dKO). **F**. Relative ^3^H-arginine uptake into Ctrl liver and L-dKO tumor tissues. Unpaired t test, ***p<0.001. n=8 (Ctrl) and 8 (L-dKO). **G**. Number of tumors per liver of L-dKO mice fed with diets containing standard concentration of arginine (100%), 10%, or 1% of arginine for 8-20 weeks of age. One-way ANOVA, ***p<0.001, ****p<0.0001. n=9 (100% arginine), 6 (10% arginine), and 9 (1% arginine). **H**. Arginine content in Ctrl liver and L-dKO non-tumor (NT) and tumor (T) tissues of mice fed with arginine-modified diets. One-way ANOVA, *p<0.05, **p<0.01, ****p<0.0001. n=6 (Ctrl, 100% arginine), 8 (Ctrl, 10% arginine), 6 (Ctrl, 1% arginine), 8 (L-dKO NT, 100% arginine), 7 (L-dKO T, 100% arginine), 6 (L-dKO NT, 10% arginine), 3 (L-dKO T, 10% arginine), 9 (L-dKO NT, 1% arginine), and 2 (L-dKO T, 1% arginine).

Arginine is a highly versatile amino acid. Besides its role as a building block in protein synthesis, it is a precursor for polyamines, creatine, and nitric oxide. Furthermore, there is interconversion between arginine, proline and glutamate, and arginine can activate mTORC1 (*8*). MPWEA yielded several terms related to arginine metabolism (arginine and proline metabolism, urea cycle, spermidine and spermine biosynthesis) among pathways up-or down-regulated in liver tumors (**table S1**). Transcriptomic and proteomic analyses of L-dKO tumors (*3*) revealed broad deregulation of arginine metabolism (**Fig. 1D**). Consistent with previous reports, we observed suppression of the urea cycle (*4, 5*) which produces arginine in the process of detoxifying excess ammonium. Decreased expression of the arginine synthesizing enzymes carbamoyl phosphate synthetase 1 (CPS1), ornithine transcarbamylase (OTC), argininosuccinate synthetase 1 (ASS1), and argininosuccinate lyase (ASL) was confirmed by immunoblotting (**Fig. 1E**). Accordingly, the urea cycle intermediates ornithine and citrulline were decreased in liver tumors (**fig. S1D**). Suppression of the urea cycle makes liver cancer cells dependent on extracellular arginine (*4, 8*). We observed up-regulation of several transporters of the solute carrier 7A family (SLC7A1, SLC7A3, SLC7A4, SLC7A6, SLC7A7, and SLC7A9) which mediate arginine uptake (*9*) (**Fig. 1, D** and **E**). We established an *ex vivo* arginine uptake assay and confirmed that arginine uptake is indeed increased in liver tumors (**Fig. 1F**). Thus, L-dKO tumors up-regulate arginine uptake to compensate for down-regulation of arginine synthesis.

Next, we investigated if high levels of arginine are critical for liver tumor development. We fed L-dKO and control mice diets that contained only 10% or 1% arginine compared to the standard diet (100% arginine), from 8 to 20 weeks of age. Control mice were not affected by the arginine-restricted diets, as assessed by liver-to-body weight ratio (**fig. S2B**). L-dKO mice displayed characteristic hepatomegaly even upon arginine restriction, again suggesting that the decreased arginine in the diets was not limiting for growth, (**fig. S2, A, B** and **C**). However, the arginine-restricted diets significantly reduced tumor burden (**Fig. 1G** and **fig. S2A**). Thus, high levels of cellular arginine are critical for the development of liver tumors.

We note that arginine levels were higher also in non-tumor liver tissue of L-dKO mice, compared to normal liver tissue of control mice, in normal diet conditions. This is likely due to the fact that at 20 weeks of age L-dKO mice develop not only multiple macroscopically visible tumors (**fig. S2A** and **Fig. 1G**) but also numerous microscopic tumors in an overall damaged liver (**fig. S2A**). This technically limits the isolation of “clean” non-tumor tissue. However, arginine levels were significantly lower in non-tumor liver tissue of L-dKO mice upon dietary arginine restriction, but unaffected (i.e., still elevated) in all tumors (**Fig. 1H**). This strict correlation of high arginine levels in tumors again suggests that high levels of cellular arginine are critical for tumorigenicity.

### Loss of ARG1 and AGMAT preserves oncogenic arginine levels

As mentioned above, arginine is a precursor for polyamines which are present in high (up to millimolar) concentrations in the cell (*10*). Thus, conversion of arginine to polyamines consumes a large amount of arginine. Polyamines, of which the major species are putrescine, spermidine and spermine, are essential for cell growth and elevated levels of polyamines have been reported in various cancer types (*10*). We detected altered expression of polyamine-related enzymes in the tumors of L-dKO mice (**Fig. 1D**). Interestingly, arginase 1 (ARG1) and agmatinase (AGMAT), which catalyze arginine-to-polyamine conversion via two distinct pathways, were down-regulated in L-dKO tumors (**Fig. 1D** and **Fig. 2A**). ARG1 cleaves arginine to produce urea and ornithine in the last step of the urea cycle. Ornithine decarboxylase (ODC) then decarboxylates ornithine to produce putrescine. In a parallel, less understood pathway, arginine is decarboxylated to produce agmatine which is then converted to putrescine by agmatinase (AGMAT). Although the conversion of arginine to agmatine has been demonstrated, there is uncertainty concerning the identity of the arginine decarboxylase that performs this step (*11*). The diamine putrescine is the precursor of the polyamines spermidine and spermine. Spermidine and spermine are synthesized sequentially by spermidine synthase (SRM) and spermine synthase (SMS), which were up-regulated in L-dKO tumors (**Fig. 1D** and **Fig. 2A**). Expression of other enzymes in polyamine metabolism were unchanged on the protein level (**Fig. 2A**). These include ODC, spermine oxidase (SMOX; converts spermine back to spermidine), spermidine/spermine N-acetyltransferase 1 (SAT1), and polyamine oxidase (PAOX; oxidizes acetylated spermine or spermidine back to spermidine or putrescine, respectively).

**Figure 2.**
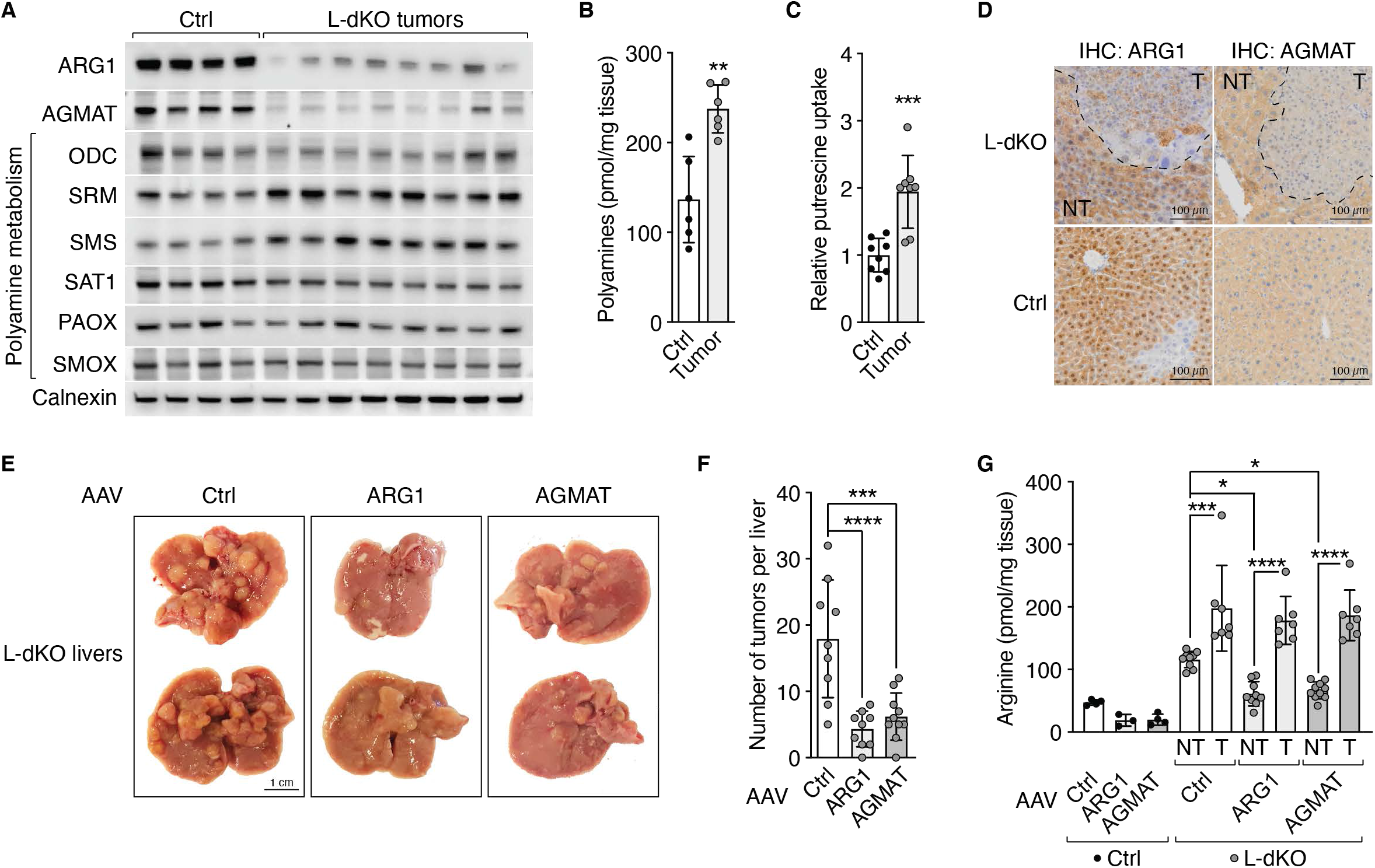
Loss of ARG1 and AGMAT enhances liver tumor formation. **A**. Immunoblot analyses of arginine-to-polyamine converting enzymes (ARG1, AGMAT) and polyamine metabolizing enzymes (ODC, SRM, SMS, SAT1, PAOX, SMOX) in Ctrl liver and L-dKO tumor tissues. Calnexin serves as loading control (same samples were used as in **Fig. 1E**). n=4 (Ctrl) and 8 (L-dKO). **B**. Total polyamine content in Ctrl liver and L-dKO tumor tissues. Unpaired t test, **p<0.01. n=6 (Ctrl) and 6 (L-dKO). **C**. Relative ^3^H-putrescine uptake into Ctrl liver and L-dKO tumor tissues. Unpaired t test, ***p<0.001. n=8 (Ctrl) and 8 (L-dKO). **D**. Immunohistochemistry analyses of Ctrl and L-dKO liver tissues stained for ARG1 or AGMAT proteins, respectively. NT (adjacent non-tumor tissue), T (tumor). **E**. Representative images of livers from L-dKO mice infected with AAV-Ctrl, AAV-ARG1, or AAV-AGMAT. **F**. Number of tumors per liver of L-dKO mice infected with AAV-Ctrl, AAV-ARG1, or AAV-AGMAT. One-way ANOVA, ***p<0.001, ****p<0.0001. n=9 (AAV-Ctrl), 9 (AAV-ARG1), and 10 (AAV-AGMAT). **G**. Arginine content in Ctrl liver and L-dKO non-tumor (NT) and tumor (T) tissues of mice infected with AAV-Ctrl, AAV-ARG1, or AAV-AGMAT. One-way ANOVA, *p<0.05. n=4 (Ctrl, AAV-Ctrl), 3 (Ctrl, AAV-ARG1), 4 (Ctrl, AAV-AGMAT), 9 (L-dKO NT, AAV-Ctrl), 7 (L-dKO T, AAV-Ctrl), 9 (L-dKO NT, AAV-ARG1), 7 (L-dKO T, AAV-ARG1), 10 (L-dKO NT, AAV-AGMAT), and 7 (L-dKO T, AAV-AGMAT).

How do the observed changes in polyamine biosynthesis enzymes affect polyamines levels in L-dKO tumors? Total polyamine levels were increased in tumors (**Fig. 2B**). Our untargeted metabolomics revealed that ions corresponding to several polyamine species were elevated, including putrescine, spermidine and the acetylated forms of spermidine and spermine (**fig. S3A**). This is consistent with reports on other cancers (*12-15*).

Our observation that both ARG1 and AGMAT are down-regulated in L-dKO tumors (**Fig. 1D** and **Fig. 2A**) raised the question, how do the tumors accumulate polyamines? Arginine can also be cleaved by mitochondrial arginase 2 (ARG2) (*7*). However, an arginine-restricted diet did not decrease polyamine levels in L-dKO non-tumor or tumor tissues (**fig. S3B**), suggesting that arginine is not the only source of polyamines in liver tumors. This is consistent with a previous report in which genetic deletion of ARG1 and/or ARG2 did not perturb polyamine homeostasis in livers of young mice (*16*). We speculated that liver tumors increase polyamine uptake. Indeed, we observed increased putrescine uptake in liver tumors (**Fig. 2C**). Thus, the intracellular pools of polyamines and arginine are uncoupled, indicating that tumors do not accumulate arginine to produce polyamines.

Why are ARG1 and AGMAT down-regulated in tumor cells? We hypothesized that loss of ARG1 and AGMAT, i.e. reducing arginine consumption, preserves the high levels of arginine which we found are critical for liver tumor development. Investigating this hypothesis, we first confirmed that loss of ARG1 and AGMAT expression is confined to tumors, by immunohistochemistry (IHC) (**Fig. 2D**). Next, to determine if expression of ARG1 and AGMAT declines early in tumor development, we performed IHC on livers of 12-and 16-week-old L-dKO animals. Of note, 12 weeks is the earliest timepoint at which defined tumors can be confirmed. Interestingly, expression of both ARG1 and AGMAT was already decreased in tumors of 12-and 16-week-old L-dKO mice (**fig. S4A**). Thus, down-regulation of ARG1 and AGMAT appears to be an early, critical event in liver tumorigenesis, possibly to preserve high levels of arginine. To test this, we injected 8-week-old L-dKO and control mice with a hepatocyte-specific adeno-associated virus (AAV) (*17*) expressing ARG1 (AAV-ARG1) or AGMAT (AAV-AGMAT). Similar to our dietary arginine restriction experiments, all AAV-infected L-dKO mice developed hepatomegaly (**Fig. 2E** and **fig. S4C**), but L-dKO mice injected with AAV-ARG1 or AAV-AGMAT developed significantly fewer tumors per liver (**Fig. 2, E** and **F**) compared to mice injected with control virus. We also observed that overexpression of ARG1 or AGMAT after AAV injection was detectable only in non-tumor liver tissue of L-dKO mice, whereas the few ‘escaper’ tumors that appeared consistently expressed low levels of ARG1 and AGMAT (**fig. S4B**). Importantly, AAV-ARG1 or AAV-AGMAT injection decreased arginine levels in non-tumor tissue, but not in the few escaper tumors (**Fig. 2G**). AAV-ARG1 or AAV-AGMAT had no effect on polyamine levels in normal liver tissue of control mice, in non-tumor tissue of L-dKO mice, or on the elevated polyamine levels in L-dKO tumors (**fig. S4D**). Taken together, our results suggest that ARG1 and AGMAT are suppressed to reduce arginine consumption and thereby preserve high levels of un-metabolized arginine required for tumorigenesis.

### Arginine determines cancer cell metabolism

How does elevated arginine, due to increased uptake and decreased consumption, promote liver cancer? To answer this question, we first screened a panel of human liver cancer cell lines for loss of ARG1 and AGMAT to find an *in vitro* experimental system that phenocopies L-dKO tumors, selecting the SNU-449 cell line (**fig. S5A**). To confirm the utility of SNU-449 cells as a proxy for L-dKO tumors, we stably expressed ARG1 or AGMAT or both ARG1 and AGMAT (hereafter ARG1/AGMAT) in these cells, using a lentivirus system (**Fig. 3A**). Expression of ARG1 and/or AGMAT only mildly reduced clonogenic growth of SNU-449 cells in standard cell culture medium (**fig. S5, B and C**). However, when grown in arginine-restricted medium, ARG1 or AGMAT expression markedly reduced clonogenic growth of SNU-449 cells, while ARG1/AGMAT co-expression arrested growth (**Fig. 3, B** and **C**). Furthermore, consistent with our *in vivo* experiments (arginine-restricted diets and AAV-mediated ARG1 or AGMAT re-expression), SNU-449 cells expressing ARG1 or AGMAT displayed reduced arginine levels, and co-expression of ARG1/AGMAT further reduced arginine levels (**Fig. 3D**). Expression of ARG1 and/or AGMAT did not increase total polyamine levels (**fig. S6A**), again consistent with our *in vivo* experiments. To investigate further the hypothesis that un-metabolized arginine promotes growth of liver cancer cells, we cultured ARG1/AGMAT expressing SNU-449 cells in the absence or presence of high levels of arginine and several arginine-related metabolites. Only L-arginine, the physiologically relevant form of arginine, and no other related metabolites (such as D-arginine, canavanine, homo-, acetyl-or methyl-arginine, or metabolites up-or downstream of arginine, such as citrulline, ornithine, agmatine, urea, and creatine) restored growth of ARG1/AGMAT expressing SNU-449 cells (**fig. S6B**). In summary, SNU-449 cells faithfully phenocopy L-dKO tumors and can thus be used as an *in vitro* proxy to study the oncogenic effect of arginine.

**Figure 3.**
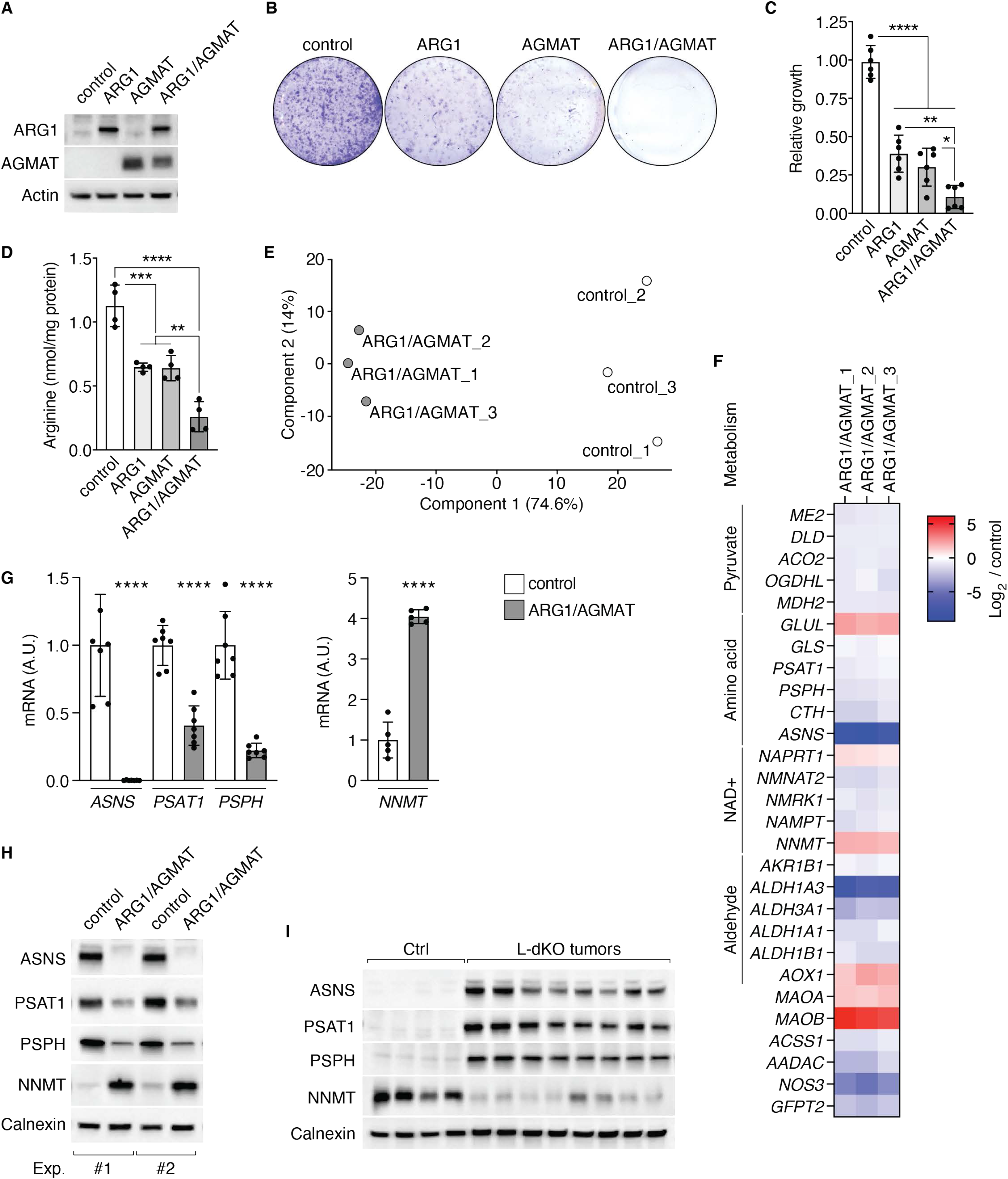
ARG1/AGMAT-controlled arginine impacts metabolic gene expression. **A**. Immunoblots of SNU-449 cells upon stable expression of ARG1 and/or AGMAT. Actin serves as loading control. **B**. Representative clonogenic growth assay of control, ARG1 and/or AGMAT expressing SNU-449 cells grown in arginine-restricted medium. **C**. Relative clonogenic growth of control, ARG1 and/or AGMAT expressing SNU-449 cells. One-way ANOVA, *p<0.05, **p<0.01, ****p<0.0001. N=6. **D**. Arginine content of control, ARG1 and/or AGMAT expressing SNU-449 cells. One-way ANOVA, **p<0.01, ***p<0.001, ****p<0.0001. N=4. **E**. PCA analysis of RNA-Seq data of control and ARG1/AGMAT expressing SNU-449 cells. **F**. Heatmap of a subset of differentially expressed metabolic genes in ARG1/AGMAT compared to control expressing SNU-449 cells (log_2_ fold-change). **G**. mRNA levels of *ASNS, PSAT1, PSPH*, and *NNMT* in control and ARG1/AGMAT expressing SNU-449 cells. Student’s t-test, ****p<0.0001. N=5-7. **H**. Immunoblots of ASNS, PSAT, PSPH, and NNMT from two independent experiments of control and ARG1/AGMAT expressing SNU-449 cells. Calnexin serves as loading control. **I**. Immunoblots of ASNS, PSAT, PSPH, and NNMT of Ctrl liver and L-dKO tumor tissues. Calnexin serves as loading control. n=4 (Ctrl) and 8 (L-dKO).

How does un-metabolized arginine promote growth of liver cancer cells? It has been reported that arginine impacts metabolism in cancer and immune cells. In leiomyosarcoma and melanoma cell lines, arginine starvation decreases glycolysis and enhances oxidative phosphorylation (OXPHOS) and serine synthesis (*18*). Conversely, in ASS1-negative breast cancer cell lines, arginine deprivation reduces OXPHOS, which leads to mitochondrial dysfunction (*19*). Also, in T cells, arginine has been shown to promote survival via regulation of multiple metabolic pathways, including enhanced OXHPOS and nucleotide synthesis (*20*). Interestingly, in prostate cancer cells arginine promotes expression of OXPHOS genes via epigenetic regulation (*21*). Thus, we examined whether elevated arginine promotes metabolic reprogramming of liver cancer cells by regulating metabolic gene expression. To identify genes differentially expressed in response to arginine, we performed RNA sequencing (RNA-seq) on SNU-449 cells expressing ARG1/AGMAT or control cells. RNA-seq samples from ARG1/AGMAT expressing cells separated from control cells in PCA and hierarchical clustering (**Fig. 3E** and **fig. S7A**). 1457 transcripts were differentially expressed ARG1/AGMAT versus control cells (**fig. S7B**), and PWEA (using KEGG pathways) was dominated by terms related to metabolism, including amino acid, NAD^+^, and glucose metabolism (**table S2** and **fig. S7C**). Since arginine has been reported to affect glycolysis and OXPHOS in other cancer types (*18, 19, 21*), we investigated if expression of ARG1/AGMAT impacted energy metabolism in liver cancer. Expression of most glycolytic genes was increased in ARG1/AGMAT expressing cells (**fig. S8A**). Notably, we found an increase in the level of mRNAs encoding glucose transporter 3 (*GLUT3*), lactate dehydrogenase A (*LDHA*), and the pyruvate dehydrogenase kinase 2 (*PDK2*) (**fig. S8, A, B** and **C**). Increased PDK2 expression was associated with an increase in inhibitory phosphorylation of the pyruvate dehydrogenase complex subunit PDHA1 on serine 293 (**fig. S8C**). These alterations suggested that ARG1/AGMAT expression, i.e., decreased intracellular arginine levels, increases glycolysis and lactate production and decreases OXPHOS. Indeed, ARG1/AGMAT expressing cells displayed reduced oxygen consumption and OXPHOS capacity (**fig. S8, D** and **E**) and secreted more lactate compared to control cells (**fig. S8F**). However, ARG1/AGMAT-controlled arginine levels impacted liver cancer cell metabolism beyond (central) energy metabolism (**table S2** and **fig. S7C**). Expression of ARG1/AGMAT also altered expression of genes in pyruvate, amino acid, NAD^+^, and aldehyde metabolism, among others (**Fig. 3F** and **fig. S8G**). From these altered metabolic pathways, we defined a “gene signature” which we used as a representative readout in further experiments. This gene signature included asparagine synthetase (*ASNS*), the serine biosynthesis genes phosphoserine aminotransferase 1 (*PSAT1*) and phosphoserine phosphatase (*PSPH*), the glucose transporter *GLUT3* and the NAD^+^ metabolic gene nicotinamide N-methyltransferase (*NNMT*). *ASNS, PSAT1*, and *PSPH* gene expression was decreased (**Fig. 3, F** and **G**) while *GLUT3* and *NNMT* expression was increased in ARG1/AGMAT expressing cells (**fig. S8, A** and **B** and **Fig. 3, F** and **G**). Accordingly, ASNS, PSAT1, and PSPH protein levels were decreased, while NNMT protein levels were increased, upon ARG1/AGMAT expression (**Fig. 3H**). Interestingly, in L-dKO liver tumors, in which ARG1 and AGMAT are suppressed (**Fig. 1D** and **Fig. 2A**), ASNS, PSAT1, and PSPH protein levels are increased, while NNMT levels are decreased (**Fig. 3I**). This correlation between ARG1/AGMAT status and expression of the signature genes further supports the hypothesis that ARG1/AGMAT-controlled arginine levels determine cancer metabolism.

### Arginine-dependent ASNS expression further enhances arginine uptake

Next, we sought to determine which of the differentially expressed genes are particularly important in linking arginine and cell metabolism in liver cancer. Interestingly, ASNS was the most differentially expressed gene in ARG1/AGMAT compared to control cells (**fig. S9A**). Furthermore, asparagine has been suggested to serve as an anti-solute in cancer cells to facilitate uptake of essential amino acids, including arginine (*22*). Of note, we observed upregulation of uniporters and antiporters that mediate arginine uptake (**Fig. 1D** and **E**). We therefore assessed uptake of arginine in ARG1/AGMAT-expressing SNU-449 cells in which ASNS is suppressed (**Fig. 3G** and **H**). Indeed, arginine uptake was reduced in ARG1/AGMAT expressing cells (**Fig. 4A**). Arginine uptake could be restored by pre-loading cells with asparagine, but not with glutamine (**Fig. 4A**) (*23*). Thus, elevated arginine uptake in liver tumors appears to depend on ASNS-derived asparagine. To test this, we stably re-expressed ASNS in SNU-449 cells expressing ARG1/AGMAT (**Fig. 4B**). Indeed, re-expression of ASNS was sufficient to elevate arginine uptake and restore clonogenic growth of SNU-449 cells expressing ARG1/AGMAT (**Fig. 4C** and **D**). Furthermore, elevated arginine uptake also restored expression of the signature genes. PSAT1 and PSPH expression was increased, while NNMT and GLUT3 expression was decreased upon ASNS expression in ARG1/AGMAT expressing cells (**Fig. 4E** and **F**). Taken together, we conclude that loss of ARG1 and AGMAT enhances arginine to promote arginine-dependent expression of ASNS. ASNS-derived asparagine further enhances arginine uptake, creating a positive feedback loop. Furthermore, the above suggests that high levels of arginine promote tumorigenesis, at least in part, by metabolic reprogramming.

**Figure 4.**
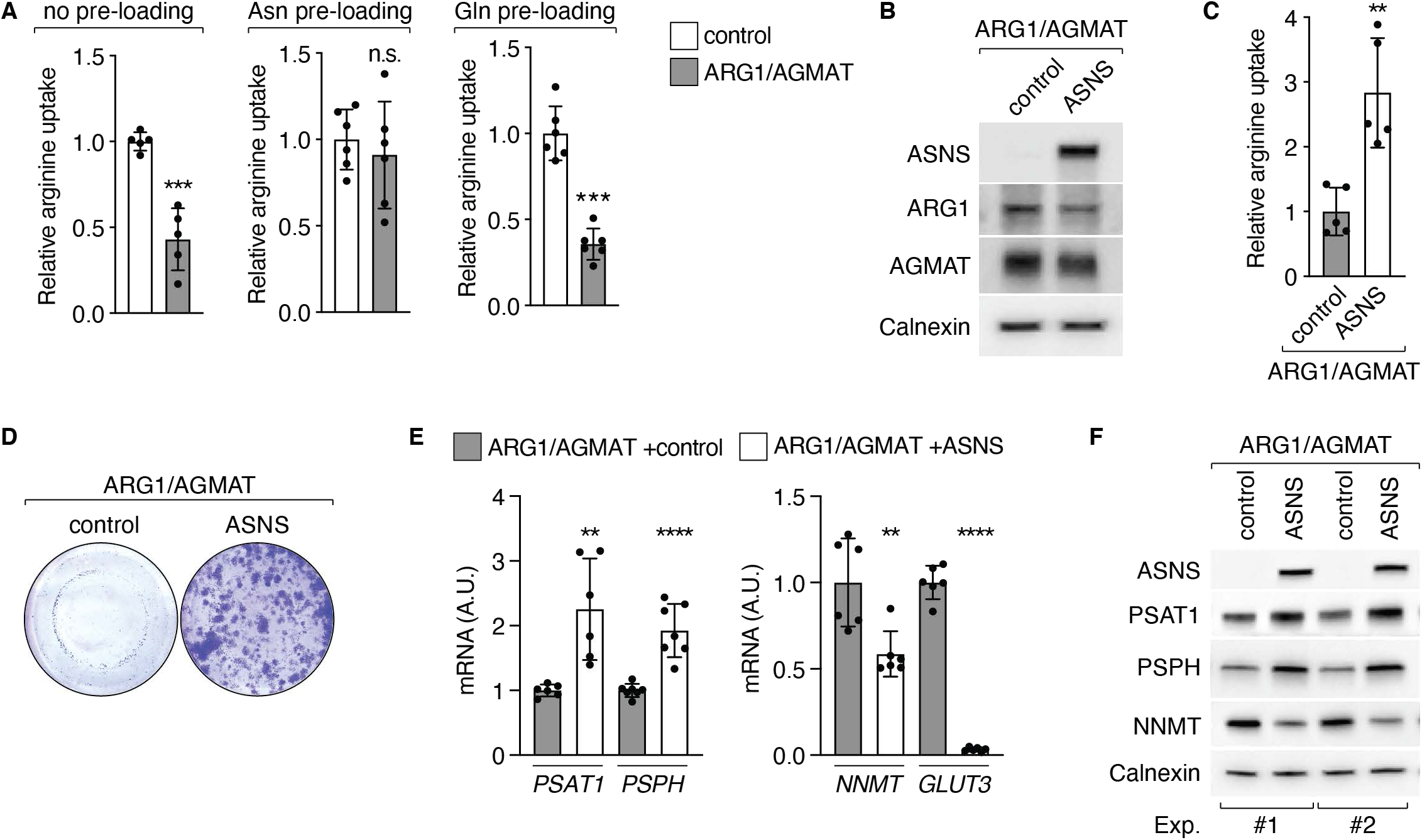
ASNS promotes arginine uptake in liver cancer. **A**. ^3^H-Arginine uptake in control and ARG1/AGMAT expressing SNU-449 cells with or without pre-loading with asparagine or glutamine. Student’s t-test, n.s.=not significant, ***p<0.001. N=5-6. **B**. Immunoblots of ARG1/AGMAT expressing SNU-449 cells upon stable expression of ASNS or control. Calnexin serves as loading control. **C**. ^3^H-Arginine uptake in control and ASNS expressing SNU-449 ARG1/AGMAT expressing cells. Student’s t-test, **p<0.01. N=5. **D**. Representative clonogenic growth assay of control and ASNS expressing SNU-449 ARG1/AGMAT expressing cells grown in arginine-restricted medium. **E**. mRNA levels of *PSAT1, PSPH, NNMT*, and *GLUT3* in control and ASNS expressing SNU-449 ARG1/AGMAT cells. Student’s t-test, **p<0.01, ****p<0.0001. N=6-7. **F**. Immunoblots of ASNS, PSAT, PSPH, and NNMT from two independent experiments of control and ASNS expressing SNU-449 ARG1/AGMAT expressing cells. Calnexin serves as loading control.

### Arginine is high in HCC patients and loss of ARG1 and AGMAT expression correlates with poor survival

Next, we analyzed proteomic and transcriptomic data obtained from biopsies of liver tumors and adjacent non-tumor tissues from liver cancer patients (*24*). We observed suppression of the urea cycle, up-regulation of several arginine transporters, deregulation of polyamine biosynthetic enzymes, and, most importantly, loss of ARG1 and AGMAT (**Fig. 5A**). These alterations were particularly pronounced in dedifferentiated, aggressive tumors (i.e., tumors clinically classified as Edmondson-Steiner high grade), which is consistent with the phenotype in our aggressive liver cancer mouse model. Immunoblotting and IHC confirmed loss of ARG1 and AGMAT and increased levels of ASNS in tumors of liver cancer patients compared to adjacent non-tumor tissue (**Fig. 5B** and **C**). Furthermore, tissue microarray analysis of more than one hundred HCC samples confirmed our finding of loss of ARG1 and AGMAT in HCC (**fig. S10A** and **Fig. 5D**). Interestingly, re-analysis of transcriptomic data from a study of early-stage HCC (*25*) revealed down-regulation of ARG1 and AGMAT, and up-regulation of ASNS (**fig. S10B**). This further supports the hypothesis that loss of ARG1 and AGMAT, and thereby, up-regulation of ASNS, are early events in HCC.

**Figure 5.**
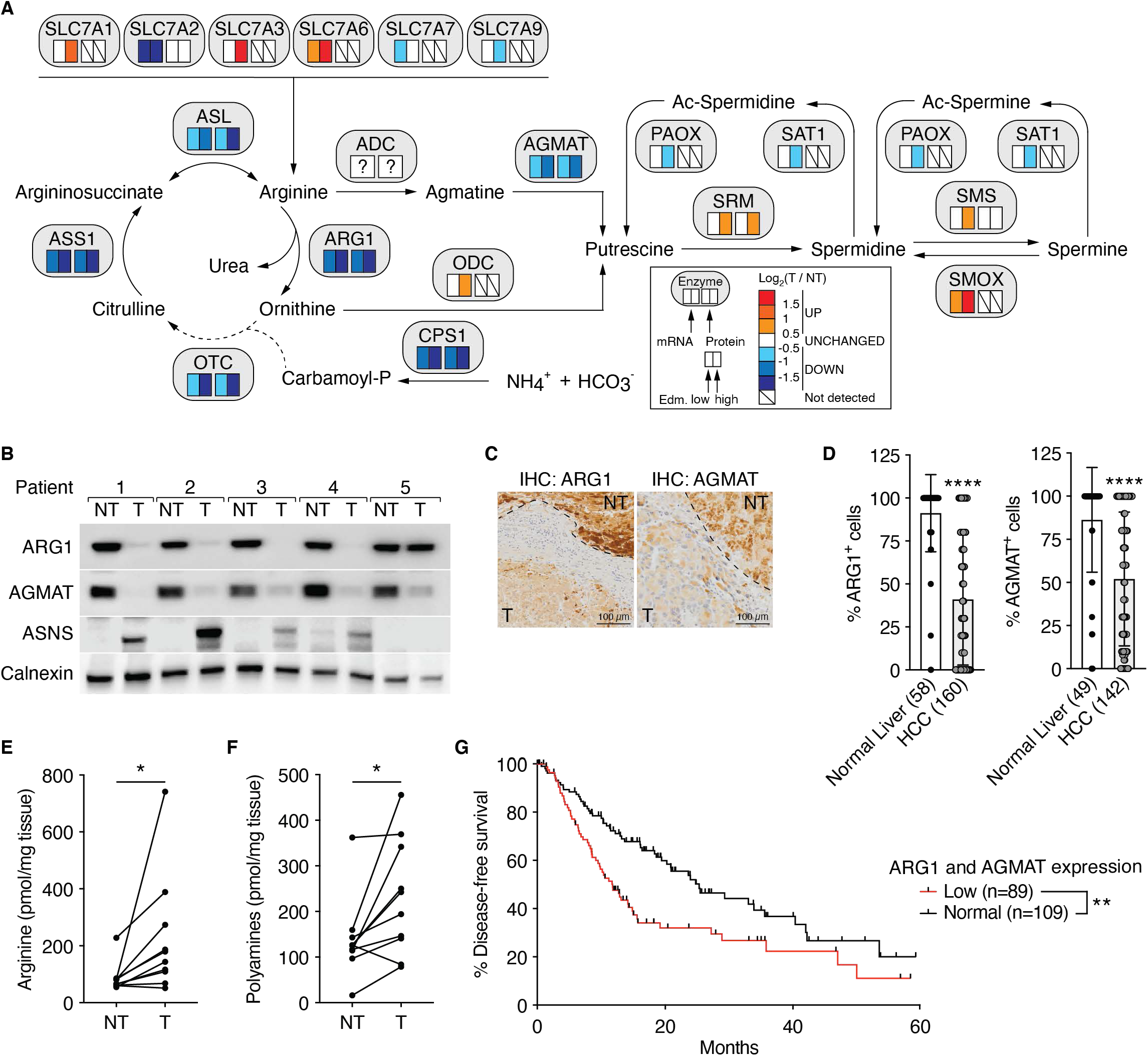
Arginine is high in human HCC patients and loss of ARG1 and AGMAT expression correlates with poor survival. **A**. Schematic representation of arginine and polyamine metabolism. Boxes below enzymes indicate changes in mRNA (left box) and protein (right box) levels in human HCC tumors (T) compared to paired non-tumor (NT) biopsies, respectively. Color coding according to level of log_2_-fold change as indicated. “?” indicates unknown identity. Tumor aggressiveness is indicated by Edmondson-Steiner grade low (Edm. low, grade I and II) and high (Edm. high, grade III and IV). n=73 (Edm. low) and 49 (Edm. high) for mRNA level, n=30 (Edm. low) and 21 (Edm. high) for protein level. **B**. Immunoblots of ARG1, AGMAT, and ASNS in paired non-tumor (NT) and tumor (T) tissues of five HCC patients. Calnexin serves as loading control. **C**. Immunohistochemistry of ARG1 and AGMAT. Non-tumor (NT) and tumor (T). **D**. Tissue microarray for ARG1 and AGMAT. Student’s t-test, ****p<0.0001. ARG1, normal liver n=58, HCC n=160; AGMAT, normal liver n=49, HCC n=142. **E**. Arginine content in paired non-tumor (NT) and tumor (T) tissues of HCC patients. Paired t-test, *p<0.05. n=10. **F**. Total polyamine content in paired non-tumor (NT) and tumor (T) tissues of HCC patients. Paired t-test, *p<0.05. n=10. **G**. Kaplan-Meier survival estimate curve for TCGA-LIHC patients ranked by expression of *ARG1* and *AGMAT*. Disease-free survival for *ARG1* and *AGMAT*: n=89 (low) and 109 (normal), log-rank test.

To assess changes in arginine metabolism-related metabolites, we performed untargeted metabolomics on 11 paired tumor and non-tumor patient biopsies. Urea cycle metabolites ornithine and citrulline were decreased, while arginine and acetylated polyamines were increased (**fig. S10C** and **D**). In addition, we biochemically validated the increase in arginine and total polyamines in an independent cohort of 10 paired tumor and non-tumor tissues (**Fig. 5E** and **F**). Finally, the importance of ARG1 and AGMAT expression in HCC patients is underscored by the finding that loss of ARG1 and/or AGMAT is associated with reduced progression-free survival, based on the TCGA liver cancer data set (**fig. S10E** and **F**, and **Fig. 5G**).

## Discussion

Alterations in the urea cycle are common in cancer (*5, 26*). It is believed that cancer cells rewire the urea cycle to enhance nitrogen usage for biomass production (*5, 27*). For example, loss of the urea cycle enzyme ASS1 in osteosarcoma increases availability of aspartate for pyrimidine synthesis (*5, 28*). However, while there has been much focus on the expression of urea cycle enzymes in cancer, there is very little known about the actual levels of arginine in these cancers. In fact, arginine levels have been evaluated only in RCC where suppression of the urea cycle results in low arginine levels (*7*). Furthermore, loss of mitochondrial ARG2 has been shown to prevent polyamine toxicity in RCC (*7*). In contrast, we describe broad deregulation of arginine metabolism which results – despite suppression of the urea cycle – in increased levels of arginine and polyamines in liver tumors. Importantly, we found that loss of ARG1 and AGMAT is a critical, and early, event in liver tumorigenesis that leads to increased levels of arginine via two mechanisms. First, loss of ARG1 and AGMAT preserves arginine by preventing arginine-to-polyamine conversion. In the absence of ARG1 and AGMAT, necessary (and high) polyamine levels are achieved via increased polyamine uptake. Thus, liver tumors seem to benefit from loss of ARG1 and AGMAT by sustaining arginine levels rather than by preventing polyamine toxicity. Second, arginine promotes expression of ASNS which utilizes aspartate to synthesize asparagine. Thus, it is tempting to speculate that loss of ASS1 in HCC provides aspartate also for ASNS. ASNS-derived asparagine further enhances arginine uptake, creating a positive feedback loop.

Several questions remain open. First, how does loss of AGMAT contribute to sustaining arginine levels? AGMAT acts downstream of ADC, i.e. on agmatine and not directly on arginine. To understand this effect, ADC first needs to be identified. One can speculate that either agmatine (which is not metabolized due to loss of AGMAT) feeds back to inhibit ADC, or that AGMAT and ADC are present in an enzyme complex and that loss of AGMAT also reduces ADC activity or expression. Second, how do liver tumors accumulate polyamines despite loss of ARG1 and AGMAT, i.e. despite suppression of arginine-to-polyamine conversion? We found elevated uptake of putrescine in liver tumors. However, polyamine transport is poorly characterized and polyamines can also be derived from glutamate, as reported in T cells (*29*). It is of interest to delineate the contribution of synthesis versus uptake, and to identify polyamine transporters in HCC. Third, and most important, how does arginine impact metabolism? We note that arginine broadly impacts metabolic gene expression, beyond ASNS. Our observation that arginine rewires metabolism at the transcriptional level suggests there is a yet-to-be determined, arginine responsive transcription factor. Further studies may reveal how arginine controls expression of ASNS and other metabolic genes to promote liver cancer.

## Acknowledgement

DM acknowledges support from the German Research Foundation (DFG). MNH acknowledges the European Research Council (MERiC), the Swiss National Science Foundation, the Sjöberg Foundation, and the Louis Jeantet Foundation.

## Figure legends

**Figure S1. Related to Figure 1**.

**A**. PCA analysis of untargeted metabolomics. n=5 (Ctrl) and 6 (T from L-dKO).

**B**. Volcano plot of the -log_10_(adjusted p-value) against the log_2_ fold-change of metabolites detected in untargeted metabolomics. Blue and red dots indicate significantly decreased and increased metabolites in tumors (T) compared to Ctrl tissues, respectively.

**C**. Arginine content in Ctrl liver and L-dKO tumor tissues, as assessed by ELISA. Unpaired t test, ***p<0.001. n=6 (Ctrl) and 6 (L-dKO).

**D**. Urea cycle metabolites in L-dKO tumors relative to Ctrl liver tissues (log_2_ ratio). Multiple t test, ***p<0.001, ****p<0.0001. n=5 (Ctrl) and 6 (L-dKO).

**Figure S2. Related to Figure 1**.

**A**. Representative images of livers from L-dKO mice fed with diets containing standard concentration of arginine (100%), 10%, or 1% of arginine for 8-20 weeks of age.

**B**. Liver-to-body weight ratio of Ctrl and L-dKO mice fed with arginine-modified diets. One-way ANOVA, n.s.=not significant. n=6 (Ctrl, 100% arginine), 9 (L-dKO, 100% arginine), 10 (Ctrl, 10% arginine), 10 (L-dKO, 10% arginine), 6 (Ctrl, 1% arginine), and 9 (L-dKO, 1% arginine).

**C**. Immunoblot analysis of mTOR signaling in Ctrl liver and L-dKO tumor tissues from mice fed with arginine-modified diets. Total S6 and total AKT serve as loading controls. n=2 (Ctrl) and 2 (L-dKO) for each diet.

**Figure S3. Related to Figure 2**.

**A**. Polyamine species in L-dKO tumors relative to Ctrl liver tissues (log_2_ ratio). Multiple t test, **p<0.01, ***p<0.001, ****p<0.0001. n=5 (Ctrl) and 6 (L-dKO).

**B**. Total polyamine content in Ctrl liver and L-dKO non-tumor (NT) and tumor (T) tissues of mice fed with arginine-modified diets. One-way ANOVA, n.s.=not significant, ***p<0.001. n=6 (Ctrl, 100% arginine), 8 (Ctrl, 10% arginine), 6 (Ctrl, 1% arginine), 8 (L-dKO NT, 100% arginine), 7 (L-dKO T, 100% arginine), 6 (L-dKO NT, 10% arginine), 3 (L-dKO T, 10% arginine), 9 (L-dKO NT, 1% arginine), and 2 (L-dKO T, 1% arginine).

**Figure S4. Related to Figure 2**.

**A**. Immunohistochemistry analyses of Ctrl and L-dKO liver tissues from 12 and 16 weeks-old mice stained for ARG1 or AGMAT proteins, respectively. NT (adjacent non-tumor tissue), T (tumor).

**B**. Immunoblot analyses of ARG1 and AGMAT levels in paired L-dKO non-tumor (NT) and tumor (T) tissues from mice infected with AAV-Ctrl, AAV-ARG1, or AAV-AGMAT. AKT serves as loading control. n=2 (AAV-Ctrl), 3 (AAV-ARG1), and 3 (AAV-AGMAT).

**C**. Liver-to-body weight ratio of Ctrl and L-dKO mice infected with AAV-Ctrl, AAV-ARG1, or AAV-AGMAT. One-way ANOVA, n.s.=not significant. n=4 (Ctrl, AAV-Ctrl), 9 (L-dKO, AAV-Ctrl), 4 (Ctrl, AAV-ARG1), 9 (L-dKO, AAV-ARG1), 4 (Ctrl, AAV-AGMAT), and 10 (L-dKO, AAV-AGMAT).

**D**. Total polyamine content in Ctrl liver and L-dKO non-tumor (NT) and tumor (Tu) tissues of mice infected with AAV-Ctrl, AAV-ARG1, or AAV-AGMAT. One-way ANOVA, n.s.=not significant, **p<0.01, ***p<0.001. n=4 (Ctrl, AAV-Ctrl), 3 (Ctrl, AAV-ARG1), 4 (Ctrl, AAV-AGMAT), 9 (L-dKO NT, AAV-Ctrl), 7 (L-dKO T, AAV-Ctrl), 9 (L-dKO NT, AAV-ARG1), 7 (L-dKO T, AAV-ARG1), 10 (L-dKO NT, AAV-AGMAT), and 7 (L-dKO T, AAV-AGMAT).

**Figure S5. Related to Figure 3**.

**A**. Immunoblot analyses of ARG1 and AGMAT levels in human liver cancer cell lines. Actin serves as loading control.

**B**. Representative clonogenic growth assay of control, ARG1 and/or AGMAT expressing SNU-449 cells grown in full (i.e. arginine-rich) medium.

**C**. Relative clonogenic growth of control, ARG1 and/or AGMAT expressing SNU-449 cells grown in full medium. One-way ANOVA, **p<0.01, ***p<0.001, ****p<0.0001. N=3.

**Figure S6. Related to Figure 3**.

**A**. Arginine content of control, ARG1 and/or AGMAT expressing SNU-449 cells. One-way ANOVA, n.s.=not significant. N=4.

**B**. Clonogenic growth of ARG1/AGMAT expressing SNU-449 cells grown in arginine-restricted medium in the presence of indicated metabolites.

**Figure S7. Related to Figure 3**.

**A**. Hierarchical clustering of RNA-Seq samples of control and ARG1/AGMAT expressing SNU-449 cells.

**B**. Volcano plot of the -log_10_(adjusted p-value) against the log_2_ fold-change of the differentially expressed genes in ARG1/AGMAT compared to control expressing SNU-449 cells. Blue and red dots indicate significantly decreased and increased gene expression, respectively.

**C**. Deregulated metabolic pathways in ARG1/AGMAT compared to control expressing SNU-449 cells after PWEA (using KEGG pathways) of differentially expressed genes from RNA-Seq.

**Figure S8. Related to Figure 3**.

**A**. Heatmap of differentially expressed glycolytic genes in ARG1/AGMAT compared to control expressing SNU-449 cells (log_2_ fold-change).

**B**. mRNA levels of *GLUT3, PDK2*, and *LDHA* in control and ARG1/AGMAT expressing SNU-449 cells. Student’s t-test, **p<0.01, ***p<0.001. N=5-7.

**C**. Immunoblots of PDK2, PDHA1 (phospho S293 and total), and LDHA from two independent experiments of control and ARG1/AGMAT expressing SNU-449 cells. Calnexin serves as loading control.

**D**. Oxygen consumption rate (OCR) of control and ARG1/AGMAT expressing SNU-449 cells. N=7.

**E**. Respiratory capacities of control and ARG1/AGMAT expressing SNU-449 cells (calculated from **D**.). Student’s t-test, ****p<0.0001.

**F**. Extracellular lactate of control and ARG1/AGMAT expressing SNU-449 cells. Student’s t-test, **p<0.01. N=4.

**G**. mRNA levels of *GLUL, GFPT2*, and *ME2* in control and ARG1/AGMAT expressing SNU-449 cells. Student’s t-test, ***p<0.001, ****p<0.0001. N=8.

**Figure S9. Related to Figure 4**.

**A**. Top ten of differentially expressed genes in ARG1/AGMAT compared to control expressing SNU-449 cells (by log_2_ fold-change, left and -log_10_(adjusted p value), right).

**Figure S10. Related to Figure 5**.

**A**. Staging of ARG1 and AGMAT immunohistochemistry (IHC) staining in tissue micro array.

**B**. mRNA expression of *ARG1, AGMAT*, and *ASNS* in early-stage HCC. log_2_ fold-change tumor (T) relative to non-tumor (NT) tissues. Multiple t test, **p<0.01, ***p<0.001, ****p<0.0001. n=35.

**C**. Urea cycle metabolites in tumors (T) relative to paired non-tumor (NT) liver tissues (log_2_ ratio). Multiple t test, *p<0.05, ****p<0.0001. n=10.

**D**. Polyamine species in tumors (T) relative to paired non-tumor (NT) liver tissues (log_2_ ratio). Multiple t test, ***p<0.001, ****p<0.0001. n=10.

**E**. Kaplan-Meier survival estimate curve for TCGA-LIHC patients ranked by expression of *ARG1*. Disease-free survival for *ARG1*: n=135 (low) and 155 (normal), log-rank test.

**F**. Kaplan-Meier survival estimate curve for TCGA-LIHC patients ranked by expression of *AGMAT*. Disease-free survival for *AGMAT*: n=136 (low) and 158 (normal), log-rank test.

