## Supplementary material for "Elevated arginine levels in liver tumors promote metabolic reprogramming and tumor growth": Materials and Methods

### Animals

Liver-specific Tsc1 and Pten double-knockout mice (L-dKO) on mixed genetic background (C57BL/6J, 129/SvJae, BALB/cJ) were as described (*1-3*). Male animals (Cre-positive (L-dKO) and Cre-negative littermates as control (Ctrl)) were used for experiments. Mice were housed at 22ºC with a 12 hour light/dark cycle with unlimited access to water and food. In all experiments, mice were fasted overnight before euthanasia by $CO_2$ inhalation. All animal experiments were performed in accordance with the federal ethical guidelines and were approved by the Kantonales Veterinäramt of Basel-Stadt.

### Arginine-modified diets

For diet experiments mice were fed with arginine-modified diets from 8-20 weeks of age. Diets were purchased from ssniff Spezialdiäten (Germany) and differed in arginine content (100% corresponding to the concentration of arginine contained in the standard diet of the animal facility (Kliba 3436)). Differences in protein/nitrogen content were balanced by increased concentrations of glycine and alanine.

### AAV administration

For AAV administration, 8-week-old Ctrl and L-dKO animals were infected with AAV8-hAAT-null (AAV-Ctrl), AAV8-hAAT-mARG1 (AAV-ARG1), or AAV8-hAAT-mAGMAT (AAV-AGMAT) via tail vein injection (2x10^12 genome copies in PBS/0.001% Pluronics F68 per mouse). AAV particles were provided by Prof. Fatima Bosch (Universitat Autònoma de Barcelona, Spain) and underlying constructs (AAV8-hAAT) were described previously (*4*).

### Cell culture

Human liver cancer cell lines SNU-182, SNU-449, HLE, PLC, Hep40, Hep3B, Huh6, Huh7, and HepG2 were gifted by Prof. Diego Calvisi (University of Greifswald, Germany), SNU-475, SNU-423, and Huh1 were gifted by Prof. Gerhard Christofori (University of Basel, Switzerland). HEK293T cells were obtained from ATCC. All cells, except Huh1, were cultured in high glucose-containing DMEM (Sigma, Cat# D5671) supplemented with 10% FBS, 2 mM glutamine, 0.1 mM non-essential amino acids (Gibco, Cat# 11140-035), and 1x penicillin/streptomycin. Huh1 cells were cultured in low glucose-containing DMEM (Sigma, Cat# D6046) supplemented with 10% FBS, 0.1 mM non-essential amino acids (Gibco, Cat# 11140-035), and 1x penicillin/streptomycin. For experiments involving modification of arginine and/or non-essential amino acids concentration in the medium, cells were cultured in DMEM lacking arginine and lysine (ThermoFisher Scientific, Cat# 88364) supplemented with 10% dialyzed FBS (ThermoFisher Scientific, Cat# 26400044), 0.798 mM lysine, 1x penicillin/streptomycin, 0.1 mM non-essential amino acids (Gibco, Cat# 11140-035) and 3.98 µM of arginine (1% compared to standard DMEM medium). Cells were incubated at 37 ºC with 5% $CO_2$ and tested for mycoplasma on regular basis.

For stable expression of ARG1 and/or AGMAT, ARG1 lentiviral Vector (pLenti-GIII-CMV-ARG1::GFP-2A-Puro, Cat# LV078906), AGMAT lentiviral Vector (pLenti-GIII-CMV-AGMAT::RFP-2A-Puro, Cat# LV071242), or Control lentiviral Vector (pLenti-CMV-GFP-2A-Puro-Blank Vector, Cat# LV590) were purchased from abmgood. For stable expression of ASNS, ASNS lentiviral Vector (pLenti-CMV-ASNS, Cat# EX-

C0302-Lv197) or Control lentiviral Vector (pLenti-CMV-Empty control vector; Cat# EX-NEG-Lv197) were purchased from genecopoeia. HEK293T cells were co-transfected with a lentiviral vector, psPAX2 (a gift from Didier Trono: Addgene plasmid # 12260) and pCMV-VSV-G (a gift from Robert Weinberg: Addgene plasmid # 8454). Supernatants containing lentiviral particles were collected 24 and 48 hours after transfection, pooled, and filtered. Filtered supernatants were used to infect cells in the presence of 10 µg/ml polybrene. Infected cells were selected with puromycin (ARG1, AGMAT, or corresponding control) or blasticidin (ASNS or corresponding control). To obtain clones with similar expression levels in single (ARG1 or AGMAT) and double (ARG1 and AGMAT) expressing cells, puromycin-selected cells were single cell-sorted by FACS for GFP (Ctrl and ARG1), RFP (AGMAT), or GFP and RFP (ARG1 and AGMAT) and expression of ARG1 and AGMAT was tested by immunoblotting.

**Clonogenic growth assays and crystal violet staining**
Low numbers of cells (500-2000 cells in 12-well plates; 1000-2500 cells in 6-well plates) were seeded in DMEM medium. Medium was exchanged the following day (to medium with modified arginine and/or asparagine concentrations as indicated) and cells were incubated for 5-14 days. To visualize colony formation ability, cells were stained with crystal violet (2% (v/v) crystal violet in 20% (v/v) methanol). Cell growth was quantified using ImageJ.

**Immunoblots**
Human and mouse liver tissues were homogenized in T-PER (ThermoFisher Scientific, Cat# 78510) supplemented with cOmplete inhibitor cocktail (Roche) and PhosSTOP (Roche) using a Polytron (PT 10-35 GT) and subsequent sonication (Hielscher UP200St). Human liver cancer cells were lysed in M-PER (ThermoFisher Scientific, Cat# 78501) supplemented with cOmplete inhibitor cocktail (Roche) and PhosSTOP (Roche). Protein concentration was determined by Pierce BCA assay (ThermoFisher Scientific, Cat# 23225), and equal amounts of protein were separated by SDS-PAGE, and transferred onto nitrocellulose membranes (GE Healthcare). Antibodies used in this study were as follows: ARG1 (GeneTex, Cat# 109242), AGMAT (Novus Biological, Cat# 1-82080), CPS1 (abcam, Cat# 129076), OTC (SantaCruz Biotech, Cat# 515791), ASS1 (SantaCruz Biotech, Cat# 365475), ASL (SantaCruz Biotech, Cat# 166787), SLC7A1 (abcam, Cat# 37588), SLC7A6 (MyBiosource, Cat# 7103267), SLC7A7 (Epigenetik, Cat# A68118-020), ODC (GeneTex, Cat# 54600), SRM (ThermoFisher Scientific, Cat# PA5-31341), SMS (SantaCruz Biotech, Cat# 376294), SAT1 (Novus Biological, Cat# 110-41622), PAOX (SantaCruz Biotech, Cat# 166185), SMOX (abcam, Cat# 213631), AKT (Cell Signaling, Cat# 4685), AKT-pS473 (Cell Signaling, Cat# 9217), Calnexin (Enzo Life Sciences, Cat# ADI-SPA-860-F), Actin (Millipore, Cat# MAB1501), ASNS (GeneTex, Cat# 30068), PSAT1 (GeneTex, Cat# 633629), PSPH (GeneTex, Cat# 33442), NNMT (abcam, Cat# 119758), S6-pS240,244 (Cell Signaling, Cat# 5364), S6 (Cell Signaling, Cat# 2217), PDK2 (SantaCruz Biotech, Cat# 517284), PDHA1-pS293 (abcam, Cat# 92696), PDHA1 (SantaCruz Biotech, Cat# 377092), LDHA (SantaCruz Biotech, Cat# 137243).

**RNA isolation and quantitative reverse transcription PCR**
Total RNA was isolated with RNeasy Kit (QIAGEN). RNA was reverse transcribed using iScript cDNA Synthesis Kit (Bio-Rad). Quantitative real-time PCR analysis was

performed using Fast SYBR Green (Applied Biosystems) and q³TOWER (Analytik Jena). Relative expression levels were determined by normalizing each Ct value to *Actin* using the ΔCt method. For each gene at least three independent biological replicates were used. The primer pairs were as follows:

| | |
|---|---|
| ASNS_forward | GGAAGACAGCCCCGATTTACT |
| ASNS_reverse | AGCACGAACTGTTGTAATGTCA |
| PDK2_forward | ATGAAAGAGATCAACCTGCTTCC |
| PDK2_reverse | GGCTCTGGACATACCAGCTC |
| LDHA_forward | TTGACCTACGTGGCTTGGAAG |
| LDHA_reverse | GGTAACGGAATCGGGCTGAAT |
| GLUT3_forward | GCTGGGCATCGTTGTTGGA |
| GLUT3_reverse | GCACTTTGTAGGATAGCAGGAAG |
| PSAT1_forward | TGCCGCACTCAGTGTTGTTAG |
| PSAT1_reverse | GCAATTCCCGCACAAGATTCT |
| PSPH_forward | GAGGACGCGGTGTCAGAAAT |
| PSPH_reverse | GGTTGCTCTGCTATGAGTCTCT |
| βActin_forward | CACCATTGGCAATGAGCGGTTC |
| βActin_reverse | AGGTCTTTGCGGATGTCCACGT |
| NNMT_forward | ATATTCTGCCTAGACGGTGTGA |
| NNMT_reverse | TCAGTGACGACGATCTCCTTAAA |
| GFPT2_forward | AGACACACTTCGGCATTGC |
| GFPT2_reverse | TTGGCGATGGTCTCTGTATCT |
| ME2_forward | ATGTTGTCCCGGTTAAGAGTAGT |
| ME2_reverse | ACCAAGCATTTGTCGTTCTTGT |
| GLUL_forward | AAGAGTTGCCTGAGTGGAATTTC |
| GLUL_reverse | AGCTTGTTAGGGTCCTTACGG |

**Arginine ELISA**
Arginine levels in mouse or human tissue homogenates or cellular lysates were measured by L-arginine ELISA kit (MyBiosource, Cat# MBS728648-96) according to the manufacturer's instructions.

**Total polyamine measurement**
Tissue or cellular lysate total polyamine levels were measured by the fluorometric Total Polyamine Assay Kit (BioVision, Cat# K475) according to the manufacturer's instructions.

**$^{3}$H-arginine and $^{3}$H-putrescine uptake**
For *ex vivo* uptake, freshly isolated Ctrl liver or L-dKO tumor tissues were incubated in 200 µl DMEM lacking arginine and 1 µCi L-[2,3,4-$^{3}$H]-Arginine (American Radiolabeled Chemicals, Cat# 1421) or 0.5 µCi Putrescine [2,3-$^{3}$H(N)] dihydrochloride (American Radiolabeled Chemicals, Cat# 0279) for 30 min at 37°C, washed twice with cold PBS and lysed with SOLVABLE (Perkin Elmer, Cat# 6NE9100). For *in vitro* uptake, cells were cultured in DMEM containing 1% arginine (compared to normal DMEM medium) for 24 hours before addition of 1 µCi L-[2,3,4-

$^{3}H$]-Arginine for 60 min at 37°C. For pre-loading of cells, 100 µM asparagine or glutamine were added to the media 30 min prior to the addition of labeled arginine. Cells were washed twice with cold PBS and lysed with 1 M HCl. Intracellular $^{3}H$-arginine or $^{3}H$-putrescine was measured with a scintillation counter.

**Oxygen consumption and lactate measurement**

Measurements were performed with an XF96 Extracellular Flux Analyzer (Seahorse Bioscience of Agilent) following manufacturer instructions. 6,500 cells were seeded into 96-well culture plate (Seahorse Bioscience of Agilent) and measured the day after. Media was exchanged prior to the measurement. Oligomycin inhibits ATP synthase (complex V), FCCP uncouples oxygen consumption from ATP production, and rotenone and antimycin A inhibit complexes I and III, respectively. All drugs were obtained from Seahorse Bioscience of Agilent.

Lactate was measured using an Arkray Lactate Pro 2 lactate test meter with corresponding test strips. Extracellular lactate was measured in the medium with background subtraction from fresh medium.

**Patient material and ethics**

All relevant ethical regulations were followed in this study. Human tissues were obtained from patients undergoing diagnostic liver biopsy at the University Hospital Basel between 2008 and 2018. Written informed consent was obtained from all patients. The study was approved by the ethics committee of the northwestern part of Switzerland (Protocol Number EKNZ 2014-099). Ultrasound-guided needle biopsies were obtained from tumor lesion(s) and the liver parenchyma at a site distant from the tumor with a coaxial liver biopsy technique that allows taking several biopsy samples through a single biopsy needle tract as described (*5, 6*). Clinical disease staging was performed using the Barcelona Clinic Liver Cancer system (*7*). In total, 122 HCC biopsies and 115 non-tumoral tissues from 114 patients with different disease etiologies were included in the study (*6*). The ethics commission approved all the experiments with resected human tissue samples reported in this Study of Northwestern Switzerland (EKNZ, approval No. 361/12).

**Proteome of HCC patients**

Fresh liver biopsies from 49 HCC were immediately snap-frozen in liquid nitrogen and processed as previously described (*3, 8*) and used for proteomic analysis (*6)*. Human HCC biopsies were measured by sequential window acquisition of all theoretical mass spectra (SWATH), in which data-independent acquisition is coupled with spectral library match (*9*). We computed the $\log_2$-fold-changes of protein abundance between paired tumor and non-tumor tissues for downstream analysis.

**RNA-sequencing and data processing of HCC patients**

RNA-seq library prep was performed with 200 ng total RNA using the TruSeq Stranded Total RNA Library Prep Kit with Ribo-Zero Gold (Illumina) according to the manufacturer's specifications. We computed the $\log_2$-fold-changes of normalized RSEM gene counts between tumors and the matched non-tumor livers for downstream analysis. RNA-sequencing data (*6*) of the human HCCs are available at the European Genome-phenome Archive under accession EGAS00001005074.

**Metabolomics of mouse liver tissues and human liver biopsies**

Snap-frozen liver samples from Ctrl and tumor tissues from L-dKO mice were collected and weighed (Ctrl: n=5, weight mean=53.7mg, stdev=4.2 mg; L-dKO: n=6, weight mean=55mg, stdev=11.4). Sample weights did not differ significantly between the two groups (two-tailed t-test, unequal variances p-value = 0.81). Snap-frozen paired tumor and non-tumor biopsies from HCC patients were weighed and 2 µl/µg extraction buffer (acetonitrile, methanol, dd$H_2O$; 2:2:1) was added. Metabolite extraction was performed as previously described (*10*). Tissue samples were kept on dry ice and homogenized in 1 ml of 70% ethanol using a Tissue Lyser 2 (Qiagen) with a stainless steel bead at maximum speed for one minute. Metabolites were extracted from the homogenized samples by adding 7ml of 70% ethanol heated to 75°C for 2minutes and subsequently cooled in ice water. Extracts were separated from cell debris by centrifuging at 2500g at 4°C for 10minutes, dried in a SpeedDry Vacuum Concentrator (Christ), and resuspended in double-distilled water (dd$H_2O$) corresponding to the measured weight, and then diluted 1:10 in dd$H_2O$ prior to mass spectrometric analysis.

Untargeted metabolomics was performed by flow injection analysis on an Agilent 6550 quadrupole instrument time-of-flight mass mass spectrometer as described previously (*11*). The instrument was operated in positive and negative mode (separate measurements), high-resolution (4GHz) mode. The injection sequence of samples was randomized, and all samples were injected in duplicates. Mass spectrometry data were pre-processed to collapse the time dimension, centroided, and merged into a single data matrix. Based on their accurate mass and the Human Metabolome Database reference list, ions were annotated, allowing tolerance of 0.003 amu and multiple common ESI adducts for initial metabolic pathway enrichment analysis (MPWEA). For subsequent analysis, annotated ions were then filtered for $H^+$ adducts allowing tolerance of 0.001 amu.

Amino acid profiling was performed by targeted metabolomics using amino acid standards from same Ctrl liver and L-dKO tumor tissues as described above.

**Statistical analysis untargeted metabolomics data**

The raw intensity data of the untargeted metabolomics analysis was processed using Perseus software (version 1.6.5.0) (*12*). $Log_2$-transformed technical duplicates were averaged and subsequently normalized by median subtraction. Separation between conditions were visualized using Perseus's built-in principal component analysis (PCA) function. Significant deregulated metabolites were determined and visualized with the volcano plot function. Thresholds were set to FDR=0.05 and S0=0.1, respectively. Unsupervised hierarchical clustering was performed after z-scoring across all rows without grouping. Spearman correlation was used to calculate the row and column tree distances. Maximum number of clusters was set to 300, iterations to 100 and restarts to 10. MPWEA was performed using the hierarchical clustering function with binarized data based on the volcano plot. Binarization was performed by defining increased metabolites as +1, decreased metabolites as -1 and unchanged metabolites as 0 following a randomization by 0.1. The pathway annotations were extracted from Small Molecule Pathway Database (SMPDB, update 2020) and linked to the measured metabolites in a FileMaker database as semicolon separated values, which could be imported to Perseus as a categorical column.

**RNA Sequencing (RNA-Seq)**
Preparation of samples for RNA-Seq, quality control, and sequencing were performed as previously described (*13*). In brief, the QuantiFluor RNA System (Promega, #E3310) was used to quantify RNA samples fluorometrically. Quality of RNA samples was checked on the TapeStation instrument (Agilent Technologies) using the High Sensitivity RNA ScreenTape (Agilent, #5067-5579). Starting from 200 ng of total RNA, library preparation was performed using the TruSeq Stranded mRNA Library Kit (Illumina, #20020595) and the TruSeq RNA Unique Dual (UD) Indexes (Illumina, #20022371) with 15 cycles of PCR. Samples were pooled to equal molarity and quantified by fluorometry using the QuantiFluor ONE double-stranded DNA System (Promega, #E4871). Libraries were quality-checked on the Fragment Analyzer (Advanced Analytical) and sequenced paired-end 51 bases using the NovaSeq 6000 instrument (Illumina) and the S1 Flow Cell loaded at a final concentration in a Flow Lane of 400 pM and including 1% PhiX. Primary data analysis was performed with the Illumina real-time analysis (RTA) version 3.4.4.

**Statistical analysis of cell line RNA-Seq**
Normalized $Log_2$ feature counts of RNA-Seq analysis were processed using Perseus software (version 1.6.14.0) (*13)*. KEGG pathway annotations were imported using Perseus's built-in annotation tool. Separation between conditions were visualized with Perseus's PCA function. Significantly differentially expressed genes were determined and visualized with the volcano plot function. Thresholds were set to FDR=0.02 and S0=0.1, respectively. Hierarchical clustering was performed after z-scoring. Maximum number of clusters was set to 300, iterations to 100 and restarts to 10. The row and tree distances were calculated using Spearman correlation. Pathway enrichment analysis (PWEA) was performed using the hierarchical clustering function with binarized data based on the volcano plot, leaving all parameters at default values. Binarization was performed by defining deregulated as 1, and unchanged as 0 following a randomization by 0.1.

**Histopathology and immunohistochemistry**
Mouse livers were fixed in 4% (w/v) paraformaldehyde, dehydrated, embedded into paraffin, cut into sections of 4 µm, and placed on SuperFrost slides (Thermo Scientific). Immunohistochemistry was performed upon Benchmark immunohistochemistry staining system (Bond, Leica) with Bond polymer refine detection solution for DAB, using ARG1 (HIER citrate buffer pH=6, 1:2500, Genetex, GTX109242), AGMAT (HIER EDTA buffer pH=9, 1:100, Sigma, PA5-55311). Immunoreactivity was evaluated by two board-certified experienced pathologists with expertise in gastrointestinal pathology (Caner Ercan and Luigi M Terraciano).

**Kaplan-Meier survival curve**
RNA sequencing gene expression data including outcomes from 298 hepatocellular carcinomas were obtained from The Cancer Genome Atlas dataset (TCGA, Provisional) via cbioportal (www.cbioportal.org). Downregulation of ARG1 or AGMAT was defined as z-score < −0.5.

**Statistics**
The investigators were not blinded to the treatment groups. Data are shown as mean ± SD. Sample numbers are indicated in each figure legend. For mouse experiments, n represents the number of animals, and for cell culture experiments, N

indicates the number of independent experiments. To determine the statistical significance paired or unpaired two-tailed Student's t test, multiple t test, or one-way ANOVA were performed using GraphPad Prism 9 Software. A p value of less than 0.05 was considered statistically significant.
