## Supplementary figures and images for "Elevated arginine levels in liver tumors promote metabolic reprogramming and tumor growth"

### Supplementary Figures S1-S10

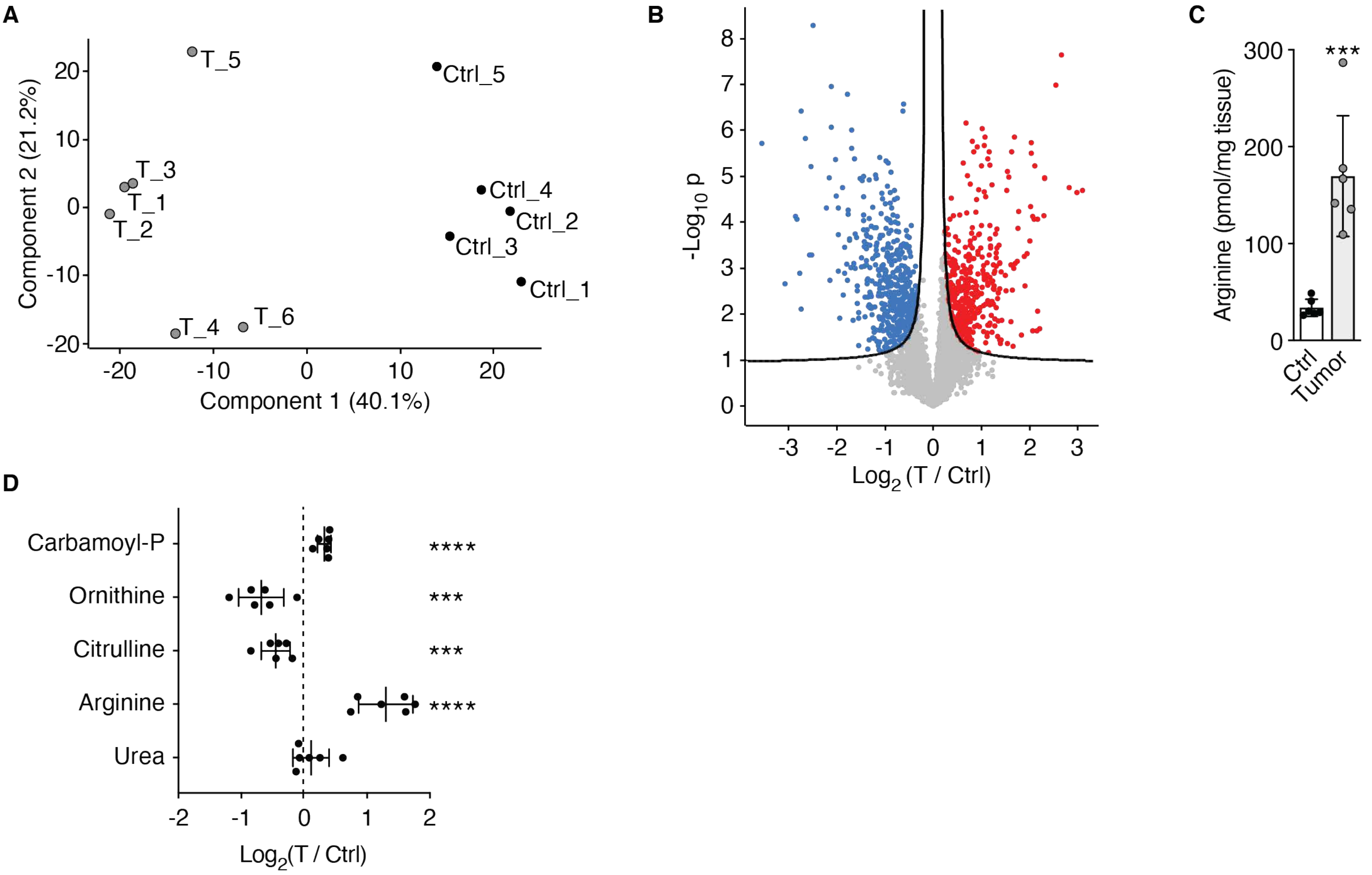

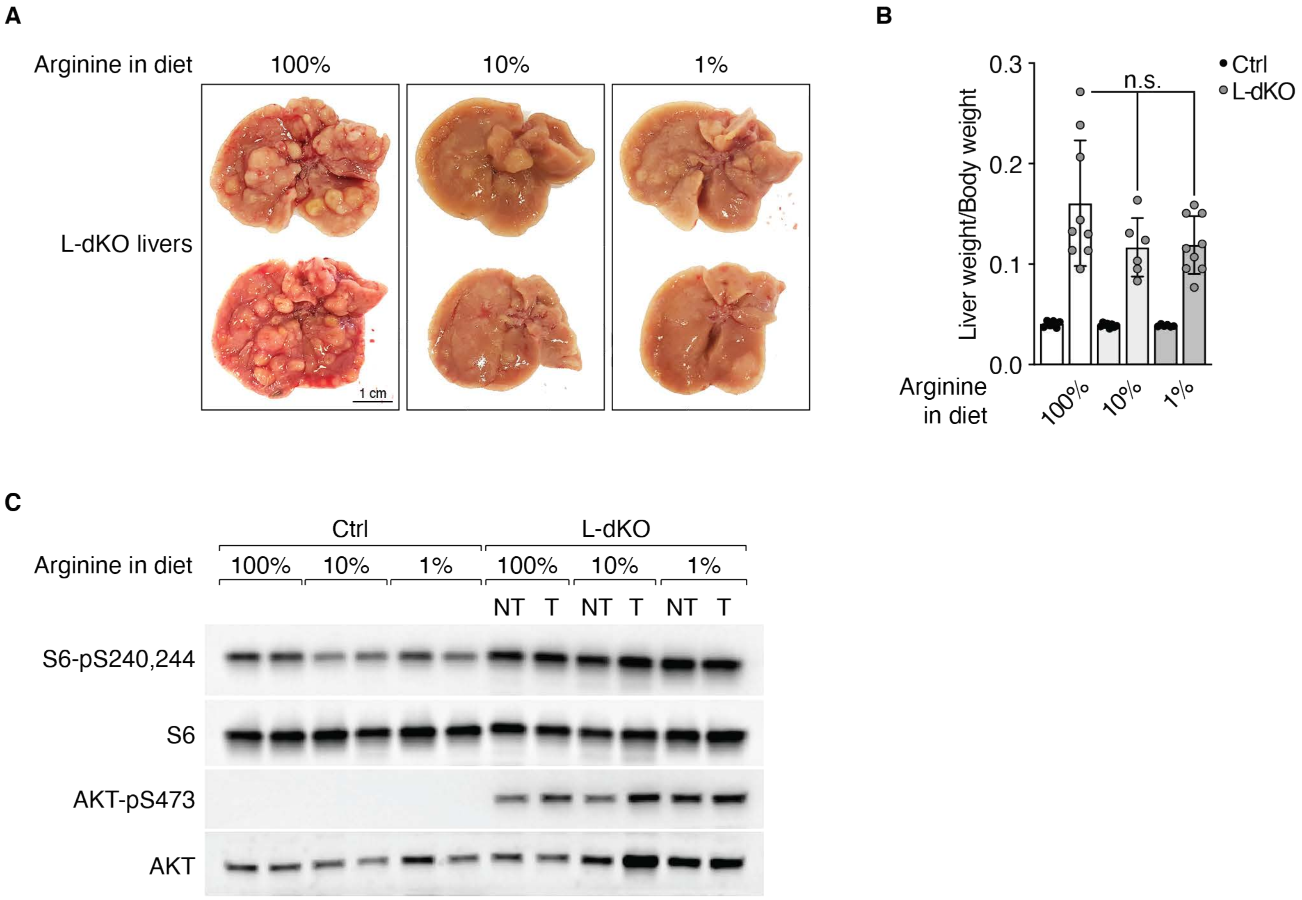

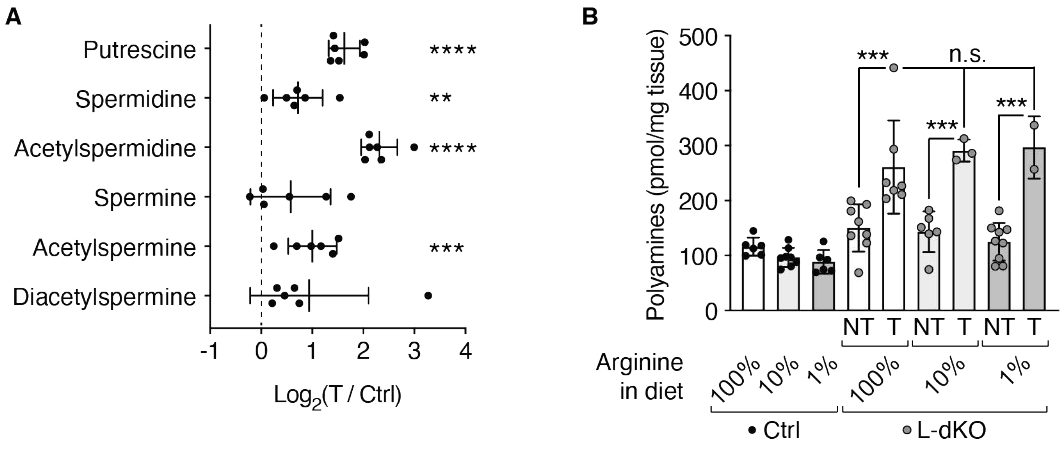

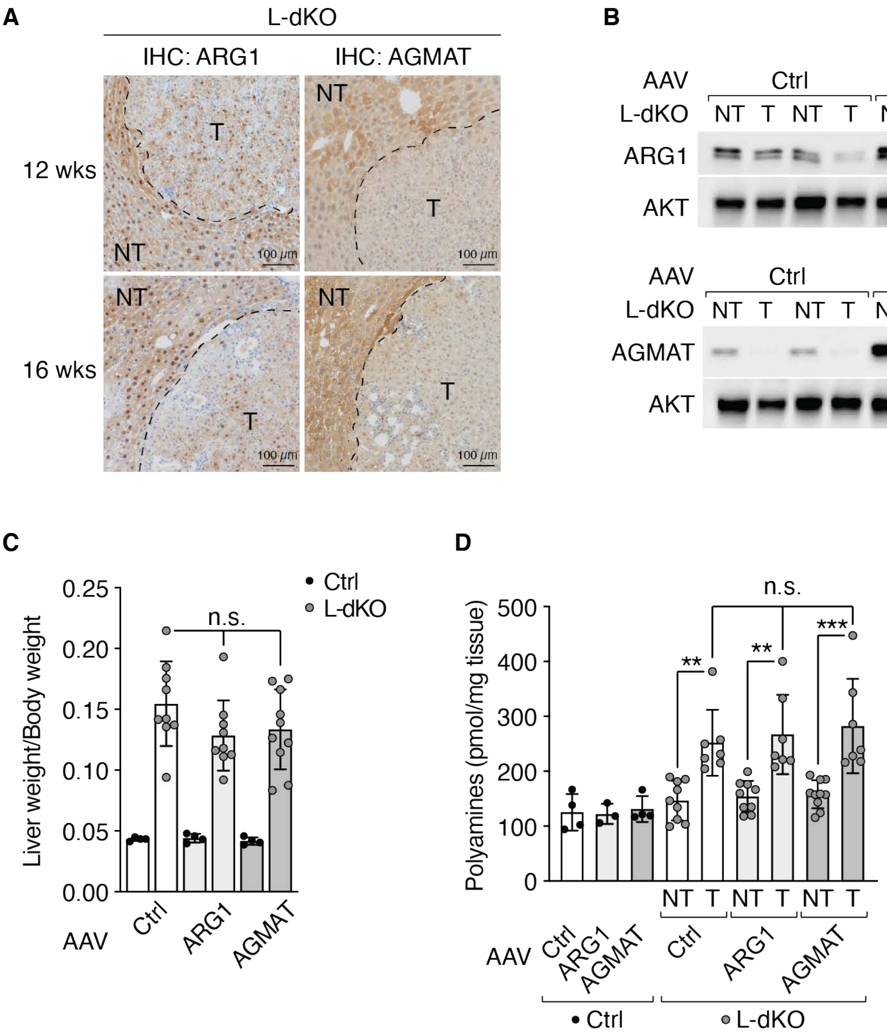

**Figure S5**

A

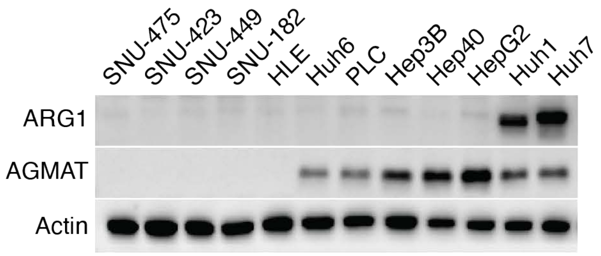


B

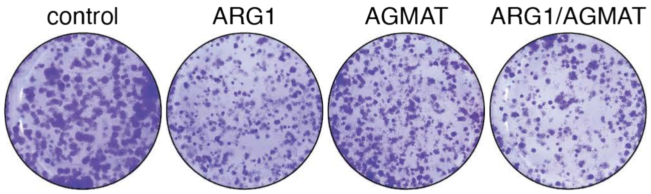


C

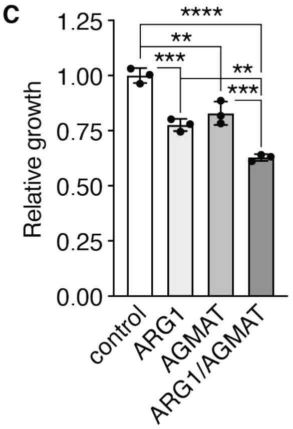

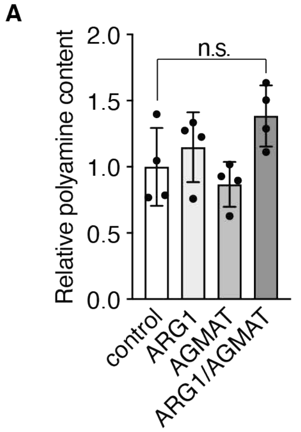

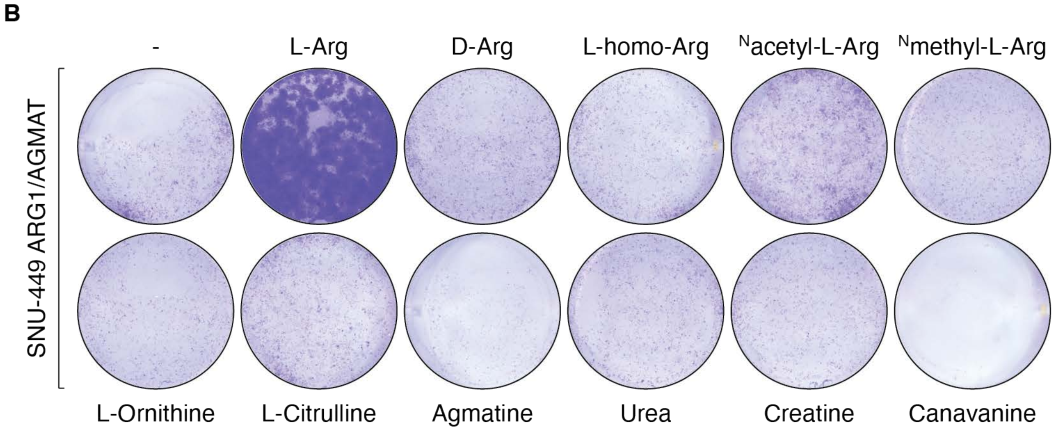

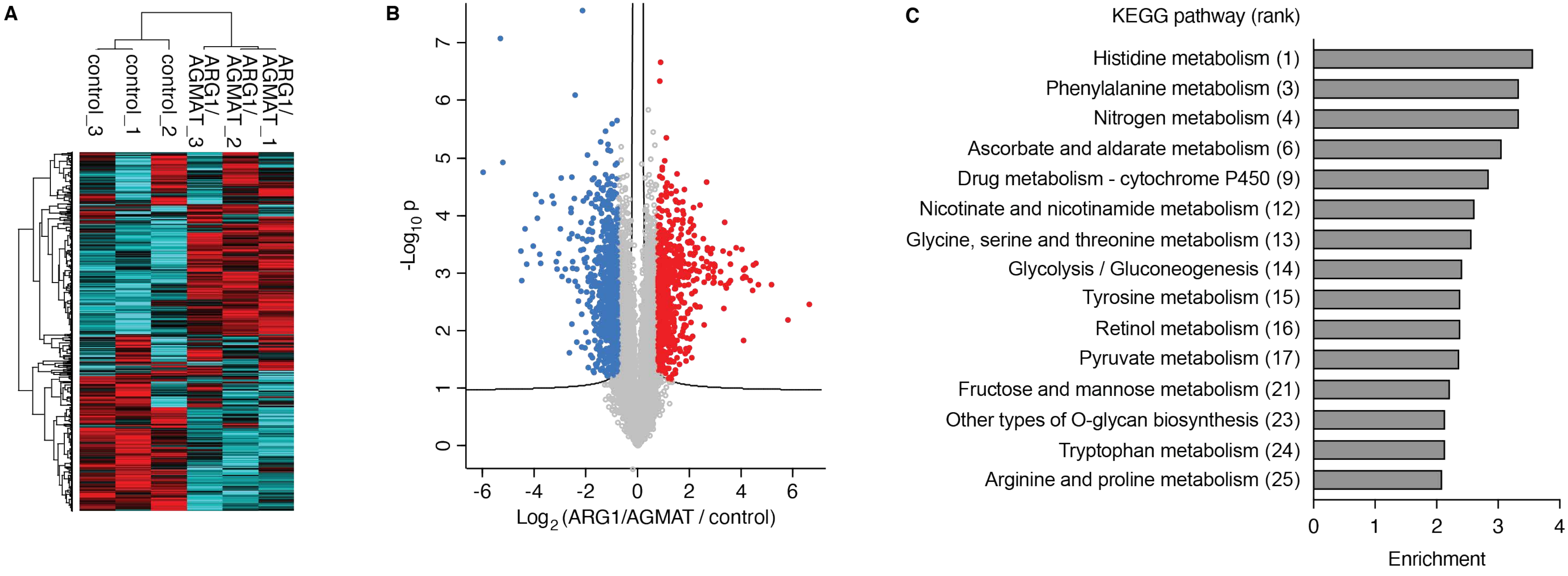

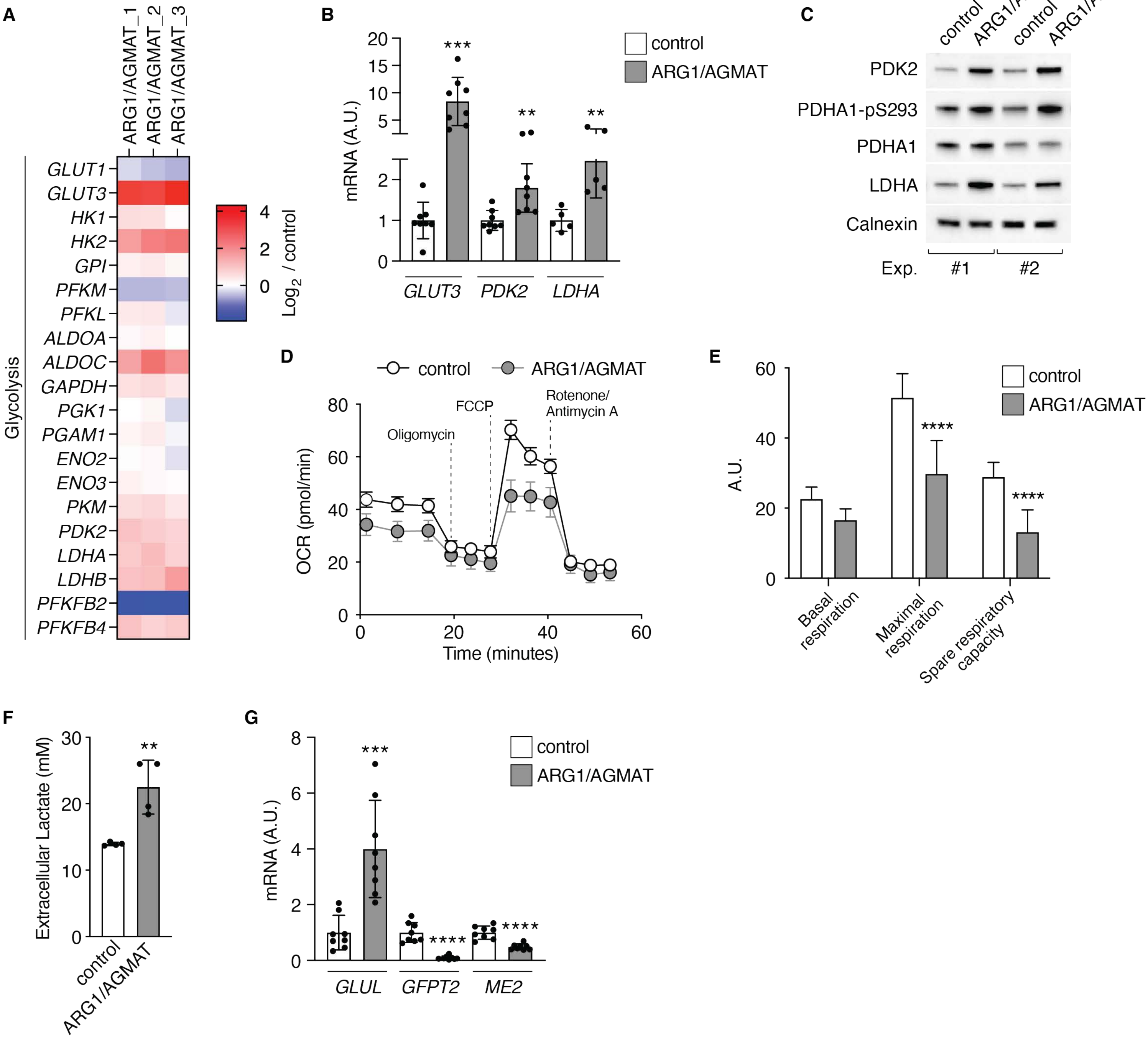

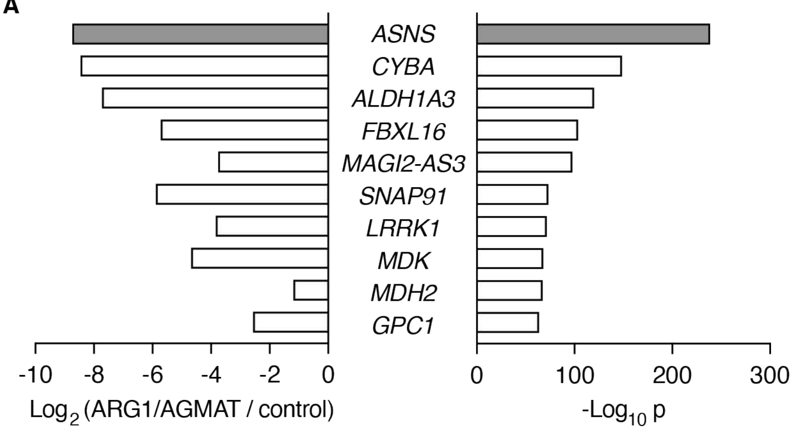

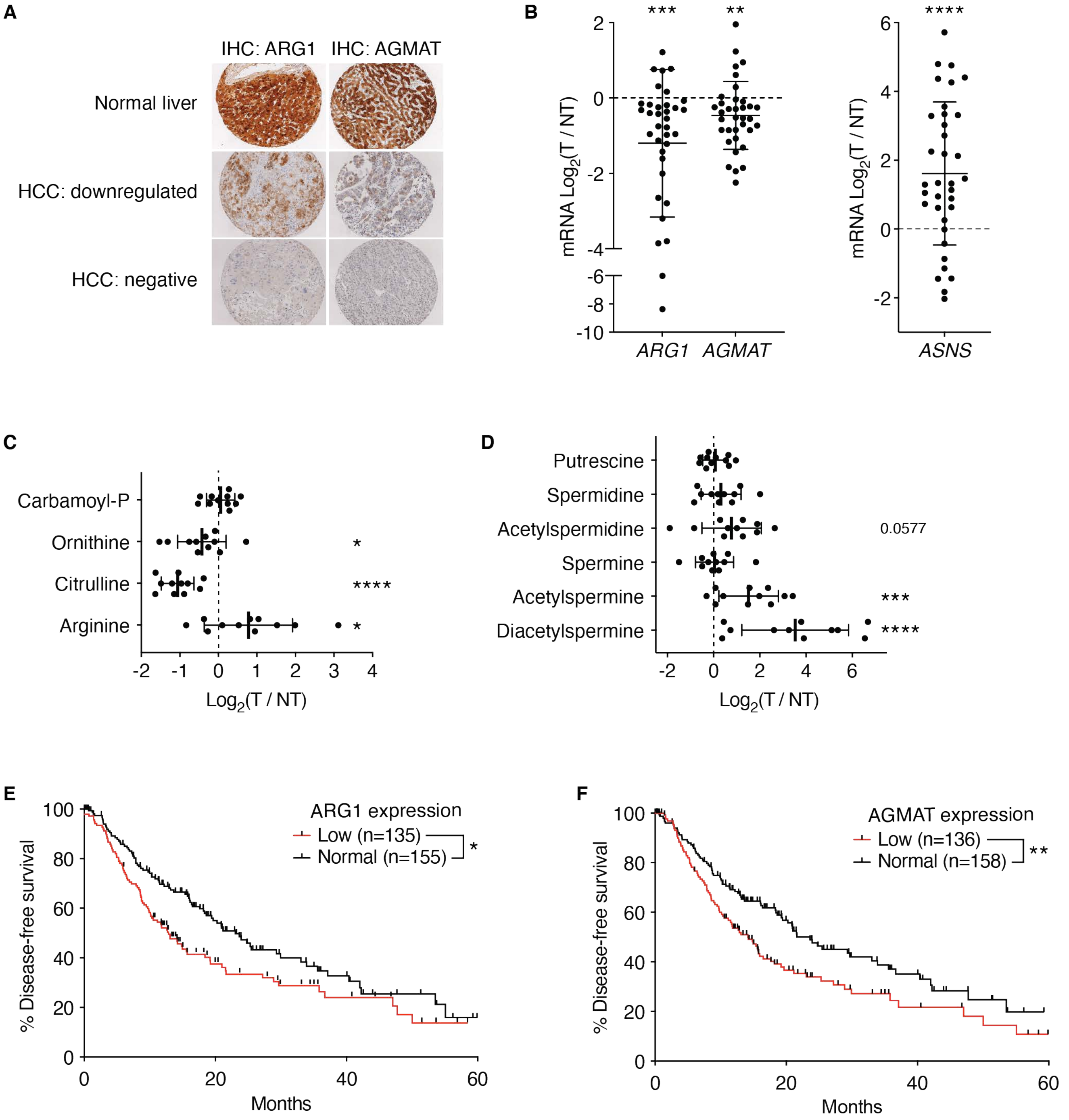
