## Supplementary material for "Elevated arginine levels in liver tumors promote metabolic reprogramming and tumor growth": Table S1 and S2

**DOWN in tumors**

| Pathway name | P value | Ben. Ho. FDR | Enrichment |
|---|---|---|---|
| Purine Metabolism | 1.1917E-08 | 1.3296E-06 | 2.4024 |
| Glycine and Serine Metabolism | 2.2859E-08 | 3.5706E-07 | 2.4493 |
| Methionine Metabolism | 3.306E-08 | 4.8716E-07 | 2.7623 |
| Ethanol Degradation | 4.9856E-08 | 6.5995E-07 | 4.4312 |
| Propanoate Metabolism | 0.000000297 | 3.3617E-06 | 2.7513 |
| Glucose-Alanine Cycle | 9.9528E-07 | 0.000010796 | 4.1895 |
| Nicotinate and Nicotinamide Metabolism | 2.5116E-06 | 0.000026507 | 2.6733 |
| Mitochondrial Beta-Oxidation of Long Chain Saturated Fatty Acids | 2.5346E-06 | 0.000026394 | 3.1127 |
| Betaine Metabolism | 3.8258E-06 | 0.000039315 | 3.0386 |
| Glutamate Metabolism | 0.00001339 | 0.00012753 | 2.1134 |
| Selenoamino Acid Metabolism | 0.000014213 | 0.00012907 | 3.3087 |
| Pantothenate and CoA Biosynthesis | 0.000016868 | 0.00015142 | 2.7743 |
| **Ammonia Recycling** | 0.000024772 | 0.00021985 | 2.3633 |
| Cysteine Metabolism | 0.000032523 | 0.00028223 | 2.6587 |
| Mitochondrial Beta-Oxidation of Short Chain Saturated Fatty Acids | 0.000047905 | 0.00037044 | 3.1783 |
| Fatty Acid Metabolism | 0.000050541 | 0.00038699 | 2.7668 |
| Mitochondrial Beta-Oxidation of Medium Chain Saturated Fatty Acids | 0.00010806 | 0.0007339 | 2.9732 |
| **Arginine and Proline Metabolism** | 0.00012938 | 0.00079566 | 1.9655 |
| Thiamine Metabolism | 0.00026214 | 0.0014836 | 3.545 |
| Phenylalanine and Tyrosine Metabolism | 0.00028716 | 0.0015794 | 2.6121 |
| Butyrate Metabolism | 0.0003108 | 0.0016512 | 2.7109 |
| Riboflavin Metabolism | 0.00047752 | 0.0023907 | 2.8885 |
| **Urea Cycle** | 0.00058493 | 0.0028731 | 2.0551 |
| Lysine Degradation | 0.00085551 | 0.0039303 | 2.5782 |
| Alanine Metabolism | 0.00086437 | 0.0038797 | 2.3633 |
| Bile Acid Biosynthesis | 0.0011039 | 0.0048165 | 2.3084 |
| Fructose and Mannose Degradation | 0.0012892 | 0.005532 | 2.3633 |
| Tryptophan Metabolism | 0.0014873 | 0.0062788 | 1.9941 |
| Valine, Leucine, and Isoleucine Degradation | 0.0015008 | 0.0058605 | 2.0429 |
| Steroidogenesis | 0.0016837 | 0.0064778 | 5.672 |
| Arachidonic Acid Metabolism | 0.0038545 | 0.013809 | 2.5524 |
| Histidine Metabolism | 0.0042485 | 0.013326 | 1.7725 |
| Malate-Aspartate Shuttle | 0.0043149 | 0.01348 | 2.7009 |
| Ketone Body Metabolism | 0.0046326 | 0.014358 | 2.9194 |
| Caffeine Metabolism | 0.0065216 | 0.019515 | 2.7572 |
| Glycolysis | 0.010386 | 0.030268 | 2.0853 |
| Nucleotide Sugars Metabolism | 0.012512 | 0.036059 | 2.0257 |
| Amino Sugar Metabolism | 0.012532 | 0.035721 | 1.8072 |
| Phenylacetate Metabolism | 0.012733 | 0.035643 | 2.127 |

**UP in tumors**

| Pathway name | P value | Ben. Ho. FDR | Enrichment |
|---|---|---|---|
| Gluconeogenesis | 7.4106E-16 | 1.1575E-13 | 4.7491 |
| Warburg Effect | 2.9539E-12 | 2.5633E-10 | 3.2478 |
| Glycolysis | 1.0241E-09 | 4.9991E-08 | 4.5373 |
| Inositol Phosphate Metabolism | 1.2804E-09 | 5.5556E-08 | 5.536 |
| Fructose and Mannose Degradation | 2.7274E-09 | 1.1211E-07 | 4.1638 |
| Citric Acid Cycle | 4.2873E-09 | 1.522E-07 | 3.543 |
| Trehalose Degradation | 6.0523E-09 | 1.5248E-07 | 8.1194 |
| Lactose Synthesis | 1.1755E-08 | 1.177E-07 | 4.0597 |
| Mitochondrial Electron Transport Chain | 1.6313E-08 | 1.6127E-07 | 4.1757 |
| Inositol Metabolism | 1.6746E-08 | 1.6348E-07 | 4.8717 |
| Starch and Sucrose Metabolism | 3.7028E-08 | 3.4842E-07 | 3.6907 |
| Transfer of Acetyl Groups into Mitochondria | 4.9291E-08 | 4.2304E-07 | 3.95 |
| Galactose Metabolism | 8.9481E-08 | 5.2943E-07 | 3.4102 |
| Amino Sugar Metabolism | 1.3404E-07 | 7.5861E-07 | 3.3433 |
| Pyruvate Metabolism | 4.1572E-07 | 0.000002042 | 2.8192 |
| Phosphatidylinositol Phosphate Metabolism | 5.0562E-07 | 2.3789E-06 | 5.2538 |
| Oxidation of Branched-Chain Fatty Acids | 5.2078E-07 | 2.4355E-06 | 3.6324 |
| Nucleotide Sugars Metabolism | 7.9127E-07 | 3.6785E-06 | 3.7117 |
| Glycerolipid Metabolism | 7.916E-07 | 3.5944E-06 | 3.3988 |
| Sphingolipid Metabolism | 1.7478E-06 | 6.2905E-06 | 3.9197 |
| **Spermidine and Spermine Biosynthesis** | 4.2685E-06 | 0.000015153 | 3.9094 |
| Pentose Phosphate Pathway | 4.4115E-06 | 0.00001559 | 3.3311 |
| Beta Oxidation of Very Long Chain Fatty Acids | 4.6262E-06 | 0.000015987 | 3.6668 |
| Sulfate/Sulfite Metabolism | 6.3599E-06 | 0.00002141 | 5.2196 |
| Phytanic Acid Peroxisomal Oxidation | 0.000010434 | 0.000034676 | 3.8973 |
| Lactose Degradation | 0.000011435 | 0.000037842 | 4.5108 |
| Thiamine Metabolism | 0.000092161 | 0.00029259 | 4.0597 |
| Phosphatidylethanolamine Biosynthesis | 0.00015691 | 0.00049214 | 3.5302 |
| Phosphatidylcholine Biosynthesis | 0.00030165 | 0.00094236 | 3.0798 |
| Steroid Biosynthesis | 0.0003781 | 0.0011143 | 3.4798 |
| Androgen and Estrogen Metabolism | 0.00057049 | 0.0016381 | 4.0597 |
| Threonine and 2-Oxobutanoate Degradation | 0.0012656 | 0.0035302 | 2.7998 |
| Alanine Metabolism | 0.0024841 | 0.006644 | 2.3198 |
| Estrone Metabolism | 0.0026112 | 0.0069602 | 2.9525 |
| Folate Metabolism | 0.0027735 | 0.0073178 | 2.7065 |
| Riboflavin Metabolism | 0.0027735 | 0.0073676 | 2.7065 |
| Bile Acid Biosynthesis | 0.0030281 | 0.0079096 | 2.2659 |
| Pyrimidine Metabolism | 0.0035178 | 0.0089201 | 1.8149 |
| Retinol Metabolism | 0.0035962 | 0.0090601 | 3.4798 |
| **Urea Cycle** | 0.0047683 | 0.011711 | 1.8828 |
| Selenoamino Acid Metabolism | 0.0057392 | 0.01392 | 2.4358 |
| Phospholipid Biosynthesis | 0.0071035 | 0.017176 | 2.3573 |
| Pantothenate and CoA Biosynthesis | 0.012915 | 0.030472 | 1.9416 |
| Propanoate Metabolism | 0.016565 | 0.038735 | 1.6966 |
| Cardiolipin Biosynthesis | 0.016954 | 0.03929 | 2.564 |
| **Arginine and Proline Metabolism** | 0.017147 | 0.038705 | 1.5274 |
| Biotin Metabolism | 0.017225 | 0.038219 | 2.8998 |

| Type | Name | P value | P value (-Log) | Enrichment | In Background list | In cluster | Ben. Ho. FDR |
|---|---|---|---|---|---|---|---|
| KEGG name | Histidine metabolism | 0.0016185 | 2.79 | 3.57 | 18 | 7 | 0.044024 |
| KEGG name | ECM-receptor interaction | 1.88E-07 | 6.73 | 3.38 | 57 | 21 | 5.11E-05 |
| KEGG name | Phenylalanine metabolism | 0.020704 | 1.68 | 3.34 | 11 | 4 | 0.13735 |
| KEGG name | Nitrogen metabolism | 0.020704 | 1.68 | 3.34 | 11 | 4 | 0.14079 |
| KEGG name | Protein digestion and absorption | 3.72E-05 | 4.43 | 3.21 | 40 | 14 | 0.005063 |
| KEGG name | Ascorbate and aldarate metabolism | 0.027679 | 1.56 | 3.06 | 12 | 4 | 0.15057 |
| KEGG name | Hematopoietic cell lineage | 0.00015893 | 3.80 | 2.98 | 40 | 13 | 0.007205 |
| KEGG name | Mineral absorption | 0.0016518 | 2.78 | 2.95 | 28 | 9 | 0.040844 |
| KEGG name | Drug metabolism - cytochrome P450 | 0.0021352 | 2.67 | 2.85 | 29 | 9 | 0.044675 |
| KEGG name | Arrhythmogenic right ventricular cardiomyopathy (ARVC) | 0.00030937 | 3.51 | 2.68 | 48 | 14 | 0.010519 |
| KEGG name | Thyroid cancer | 0.008831 | 2.05 | 2.68 | 24 | 7 | 0.1201 |
| KEGG name | Nicotinate and nicotinamide metabolism | 0.016034 | 1.79 | 2.62 | 21 | 6 | 0.12461 |
| KEGG name | Glycine, serine and threonine metabolism | 0.010933 | 1.96 | 2.57 | 25 | 7 | 0.11014 |
| KEGG name | Glycolysis / Gluconeogenesis | 0.0043647 | 2.36 | 2.42 | 38 | 10 | 0.0742 |
| KEGG name | Tyrosine metabolism | 0.023697 | 1.63 | 2.39 | 23 | 6 | 0.13714 |
| KEGG name | Retinol metabolism | 0.023697 | 1.63 | 2.39 | 23 | 6 | 0.14012 |
| KEGG name | Pyruvate metabolism | 0.010971 | 1.96 | 2.37 | 31 | 8 | 0.1029 |
| KEGG name | Malaria | 0.010971 | 1.96 | 2.37 | 31 | 8 | 0.10657 |
| KEGG name | Hypertrophic cardiomyopathy (HCM) | 0.0014879 | 2.83 | 2.29 | 56 | 14 | 0.044967 |
| KEGG name | Complement and coagulation cascades | 0.015342 | 1.81 | 2.23 | 33 | 8 | 0.12274 |
| KEGG name | Fructose and mannose metabolism | 0.022408 | 1.65 | 2.22 | 29 | 7 | 0.13852 |
| KEGG name | Axon guidance | 0.00012875 | 3.89 | 2.20 | 96 | 23 | 0.0070039 |
| KEGG name | Other types of O-glycan biosynthesis | 0.026051 | 1.58 | 2.14 | 30 | 7 | 0.14461 |
| KEGG name | Tryptophan metabolism | 0.026051 | 1.58 | 2.14 | 30 | 7 | 0.14762 |
| KEGG name | Arginine and proline metabolism | 0.011498 | 1.94 | 2.09 | 44 | 10 | 0.10425 |
| KEGG name | Amoebiasis | 0.0021708 | 2.66 | 2.07 | 71 | 16 | 0.042176 |
| KEGG name | Cytokine-cytokine receptor interaction | 0.00010456 | 3.98 | 2.04 | 126 | 28 | 0.0071102 |
| KEGG name | Metabolism of xenobiotics by cytochrome P450 | 0.043851 | 1.36 | 2.04 | 27 | 6 | 0.19879 |
| KEGG name | Rheumatoid arthritis | 0.010582 | 1.98 | 2.02 | 50 | 11 | 0.11993 |
| KEGG name | Basal cell carcinoma | 0.021338 | 1.67 | 1.97 | 42 | 9 | 0.13819 |
| KEGG name | Dilated cardiomyopathy | 0.010918 | 1.96 | 1.93 | 57 | 12 | 0.11422 |
| KEGG name | Small cell lung cancer | 0.0057452 | 2.24 | 1.91 | 72 | 15 | 0.086816 |
| KEGG name | Neuroactive ligand-receptor interaction | 0.0016673 | 2.78 | 1.89 | 102 | 21 | 0.037791 |
| KEGG name | Bile secretion | 0.034096 | 1.47 | 1.88 | 39 | 8 | 0.16561 |
| KEGG name | Epithelial cell signaling in Helicobacter pylori infection | 0.017124 | 1.77 | 1.87 | 54 | 11 | 0.12589 |
| KEGG name | Focal adhesion | 0.00028383 | 3.55 | 1.85 | 154 | 31 | 0.011029 |
| KEGG name | Ribosome | 0.0089246 | 2.05 | 1.81 | 76 | 15 | 0.1156 |
| KEGG name | PPAR signaling pathway | 0.042068 | 1.38 | 1.79 | 41 | 8 | 0.19728 |
| KEGG name | Pathways in cancer | 6.35E-05 | 4.20 | 1.71 | 257 | 48 | 0.0057561 |
| KEGG name | Lysosome | 0.0083161 | 2.08 | 1.68 | 104 | 19 | 0.11905 |
| KEGG name | Osteoclast differentiation | 0.021706 | 1.66 | 1.60 | 86 | 15 | 0.1373 |
| KEGG name | Chemokine signaling pathway | 0.01444 | 1.84 | 1.57 | 111 | 19 | 0.11902 |
| KEGG name | Leukocyte transendothelial migration | 0.032666 | 1.49 | 1.53 | 84 | 14 | 0.16454 |
| KEGG name | MAPK signaling pathway | 0.019281 | 1.71 | 1.35 | 204 | 30 | 0.13447 |
